# Most regulatory interactions are not in linkage disequilibrium

**DOI:** 10.1101/272245

**Authors:** Sean Whalen, Katherine S. Pollard

## Abstract

Linkage disequilibrium (LD) and genomic proximity are commonly used to map non-coding variants to genes, despite increasing examples of causal variants outside the LD block of the gene they regulate. We compared chromatin contacts in 22 cell types to LD across billions of pairs of loci in the human genome and found no concordance, even at genomic distances below 25 kilobases where both tend to be high. Gene expression and ontology data suggest that chromatin contacts identify regulatory variants more reliably than do LD and genomic proximity. We conclude that the genomic architectures of genetic and physical interactions are independent, with important implications for gene regulatory evolution and precision medicine.

---

Genetic variants ranging from large scale chromosomal rearrangements to single nucleotide polymorphisms (SNPs) can impact gene function by altering exonic sequence or by changing gene regulation. Recent studies estimate that 93% of disease-associated variants are in non-coding DNA [1] and 60% of causal variants map to regulatory elements [2], accounting for 79% of phenotypic variance [3]. Additionally, disease-associated variants are enriched in regulatory regions [4], especially those from tissues relevant to the phenotype [5]. Functionally annotating non-coding variants and correctly mapping causal variants to the genes and pathways they affect is critical for understanding disease mechanisms and using genetics in precision medicine [6–9].

Common practice associates non-coding variants with the closest gene promoter or promoters within the same LD block. However, regulatory variants can affect phenotypes by changing the expression of target genes up to several megabases (mb) away [10–13], well beyond their LD block (median length ≈ 1-2kb, Supplementary Table 1b). This prompted Corradin and colleagues to conclude that a gene’s regulatory program is not related to local haplotype structure [14]. Even when a GWAS SNP is in LD with a gene that has a strong biological link to the phenotype, the causal variant may be in a nearby non-coding region regulating a different gene [15, 16]. Highlighting the long range of regulatory interactions, recent work in T cells found that only 14% of 684 autoimmune variants targeted their closest gene; 86% skipped one or more intervening genes to reach their target, and 64% of variants interacted with multiple genes [17]. Thus, many non-coding variants are far away and in low LD with the promoters they regulate.

Distal non-coding variants can cause changes in gene regulation and phenotypes via three-dimensional (3D) chromatin interactions. For example, the obesity-associated FTO variant (rs1421085) was found to disrupt an ARID5B repressor motif in an enhancer for IRX3/5 during adipocyte differentiation, increasing obesity risk [10]. A second study showed a schizophrenia-associated SNP (rs1191551) regulates the expression of distal gene FOXG1 in two zones of the developing human cerebral cortex, rather than targeting the nearby gene PRKD1 [13]. Another example is a papillary thyroid cancer associated SNP (rs965513) in an LD block containing several enhancer variants that contact the promoter of FOXE1 and alter its expression [18]. In addition, mutagenesis screens identified multiple distal variants that lead to cancer drug resistance by decreasing CUL3 expression [19]. These validated causal SNPs demonstrate that regulatory variants can be located far from their target promoters in distinct LD blocks (IRX3/5 1.2mb, FOXG1 760kb, CUL3 ± 100kb, FOXE1 ± 60kb).

New understanding of the 3D genome from high-throughput chromatin capture (Hi-C) and imaging data suggests it may be common for regulatory variants and their target gene(s) to lack strong LD. For example, mammalian genomes are partitioned into regions enriched for chromatin interactions at multiple scales, including Topologically Associating Domains (TADs, median length 880kb [20]) and contact domains (sub-TADs, median length 250kb [21]). While these chromatin domains resemble the nested block patterns of LD, they have a different origin: insulating chromatin boundary elements across which relatively few chromatin interactions occur versus frequency of recombination events over generations. Furthermore, different proteins interact with DNA to mediate these processes, namely PRDM9 in the case of recombination [22] and structural proteins such as CTCF in the case of chromatin boundaries [23]. Thus, one might not expect similarity *a priori*. However, LD is high and chromatin interactions are common at genomic distances less than 25 kilobases (kb), so LD and chromatin contact maps might be correlated at this scale even though some causal SNPs regulate promoters over long genomic distances where LD is approximately zero. The correspondence between chromatin contact frequencies and LD has yet to be comprehensively evaluated on a genome-wide scale.

To address this question, we quantified the relationship between LD and chromatin contact frequency genome-wide using Hi-C data from 5 diverse cell lines [21] and promoter capture Hi-C (PCHi-C) data from 17 primary blood cell types [12], combined with SNPs from the 1000 Genomes Project [24]. In several analyses, we focus on blood cells to enable integration of B cell eQTLs [25] and blood-relevant data from the GWAS catalog [1] with high resolution chromatin interaction data in a consistent cellular context. Using multiple large scale analyses over billions of chromatin contacts and SNP pairs, we first demonstrate that LD is not a proxy for regulatory interaction. In particular, LD decays at much shorter distances than chromatin contacts and is no higher in regions with statistically significant chromatin interactions than distance matched regions with non-significant interactions. Even at genomic distances < 25kb where LD and contact frequencies are both high, their block patterns are rarely aligned. We then show that utilizing higher-order chromatin organization is essential for accurate functional assignment of non-coding variants, having implications for fine mapping of causal variants and applications to precision medicine.

## Results

To comprehensively compare the genomic architectures of LD and chromatin contacts, we generated two types of data structures from publicly available data (Supplementary Table 1). The first includes LD blocks and pairwise LD between all high-quality, bi-allelic SNPs across individuals from each of the 1000 Genomes Project superpopulations [24] (AFR: African, AMR: Ad Mixed American, EAS: East Asian, EUR: European, SAS: South Asian). The second records contact frequencies between all pairs of fragments in 22 human cell types with high-resolution Hi-C [21] or promoter capture Hi-C (PCHi-C) [12] data that measures interactions between baited promoters and promoter interacting regions (PIRs). These chromatin contact data were used to generate lists of statistically significant interacting regions and distance-matched regions with non-significant interactions for each cell type using with methods that account for expected contact frequencies and adjust for multiple testing. Significant chromatin interactions from Hi-C were computed using Juicer [26] and represent statistical enrichment of contacts over a particular choice of local background, whereas those from PCHi-C were identified using CHiCAGO [27] and indicate if a region is likely to be in the same contact domain with a promoter or not. Due to the resolution of chromatin interaction assays, we could not compare LD and 3D proximity between sites separated by less than 5kb where both values are expected to be high.

The resulting analyses span approximately 1.6 million LD blocks, 27 billion SNP pairs, and 3.1 million statistically significant chromatin interactions (Supplementary Table 1). By analyzing the genome-wide relationship between LD and chromatin contacts from multiple perspectives, we show that LD is not correlated with chromatin interactions and generally should not be used for mapping non-coding SNPs to genes and pathways.

### Chromatin interactions and LD have different genomic architectures

Both LD and chromatin contact frequency measure the strength of a relationship between pairs of genomic sites. However, these two measures differ fundamentally in their scales: Chromatin contacts span much longer distances (Figure 1). Genetic architecture forms LD blocks of median length 2kb (combined 1000 Genomes super-populations) in which a percentage of SNP pairs exceed a common threshold of *R*^2^ > 0.8 [28]. Strong LD pairs have a median distance of 13kb as they can be located in different blocks. On the other hand, physical architecture forms regions enriched for interactions at much longer scales, including contact domains (median 250kb [21]), focal interactions (median 270kb [21] or 350kb [12]), and topological domains (median 840kb). This difference is evident in most genomic loci when contact frequency from a particular cell type is plotted alongside LD from 1000 Genomes, both at the scale of TADs (Figure 2a) and within smaller contact domains (Figure 2b) where chromatin interactions are frequent but LD structure is low or limited to smaller LD blocks. Due to this difference in scale, non-coding SNPs frequently contact genes located hundreds of kb away without being in LD with those genes (Figure 2b). These distal chromatin interactions may differ across cell types, whereas LD does not (Figure 2c). A similar example is shown in Supplementary Figure 1.

To quantify the decay rates of LD versus chromatin contacts genome-wide, we analyzed all pairs of sites separated by a given genomic distance with respect to Hi-C contact frequency in ENCODE cell lines [21] versus LD in 1000 Genomes individuals. This showed that contact frequency decays with genomic distance much slower than LD both across (Figure 3, Supplementary Figure 2) and within human populations (Supplementary Figure 4). Furthermore, statistically significant chromatin interactions occur between genomic regions separated by dozens, hundreds, or even thousands of LD blocks (Supplementary Figure 6 panels a-b), while most SNP pairs with non-zero LD cross 0-2 contact domains (Supplementary Figure 6, panel c). PCHi-C data shows the same broad trends (Supplementary Figure 3). In summary, genetic and physical architectures of human chromosomes differ at all scales (Figure 1).

### Chromatin contact frequencies have low concordance with LD across genomic distances

Contact frequency and LD could still be correlated at shorter genomic distances where LD is more often non-zero. To explore this possibility, we analyzed the concordance of frequent Hi-C contacts and strong LD values (*R*^2^ > 0.8) for pairs of sites within genomic windows from 5kb to 1.2mb (Methods). Chromatin interactions and strong LD co-occur about as often as expected if there was no association between the two variables (Figure 4). Concordance is highest at short genomic distances and decays as LD approaches zero. Notably, concordance is nearly zero at scales where statistically significant chromatin interactions still occur. The maximum concordance and rate of decay differ across 1000 Genomes super-populations, with the lowest concordance in African populations that have the fastest decay of LD. These patterns are consistent with the two variables being independent and LD decaying more rapidly with genomic distance (Figure 3). Hence, there is no evidence that LD and chromatin contacts are correlated at genomic distances of 5kb or more.

**Figure 1:**
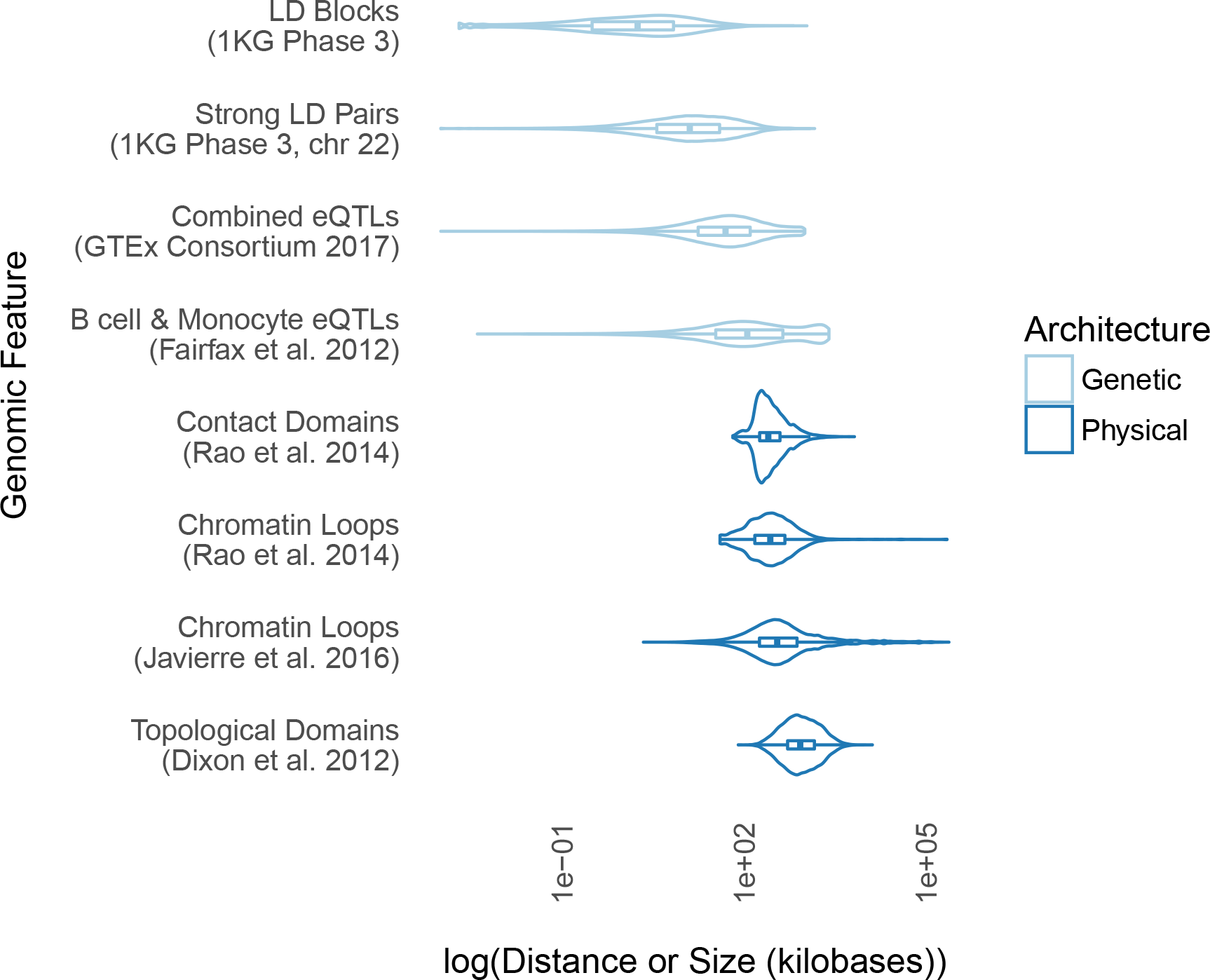
Genetic and Physical Architectures. LD blocks and strong LD pairs (*R*^2^ > 0.8) operate across tens of kb or less, while chromatin interactions and multi-scale domains of enrichment span hundreds of kb, with eQTLs roughly in between. Summaries are computed over all super-populations, tissue types, or cell types in the relevant datasets.

### LD is not elevated in significant chromatin interactions

Next we compared LD and chromatin structure focusing on statistically significant chromatin interactions, as these might harbor high LD SNPs even if less frequent chromatin contacts are rarely genetically linked. For each statistically significant and distance-matched non-significant interaction, we computed the maximum LD between pairs of SNPs occurring on opposing fragments. The ratio of interacting versus non-interacting fragment LD is often close to 1 across all super-populations and cell types (Figure 5, Supplemental Table 2), indicating no elevation of LD at interacting regions. In addition, LD is very low between non-coding regions and interacting promoters in PCHi-C data, with 2-7% of interacting fragments located within the same LD block (Table 1). Thus, LD is not an adequate proxy for high-confidence chromatin interactions, including those involving promoters.

**Figure 2:**
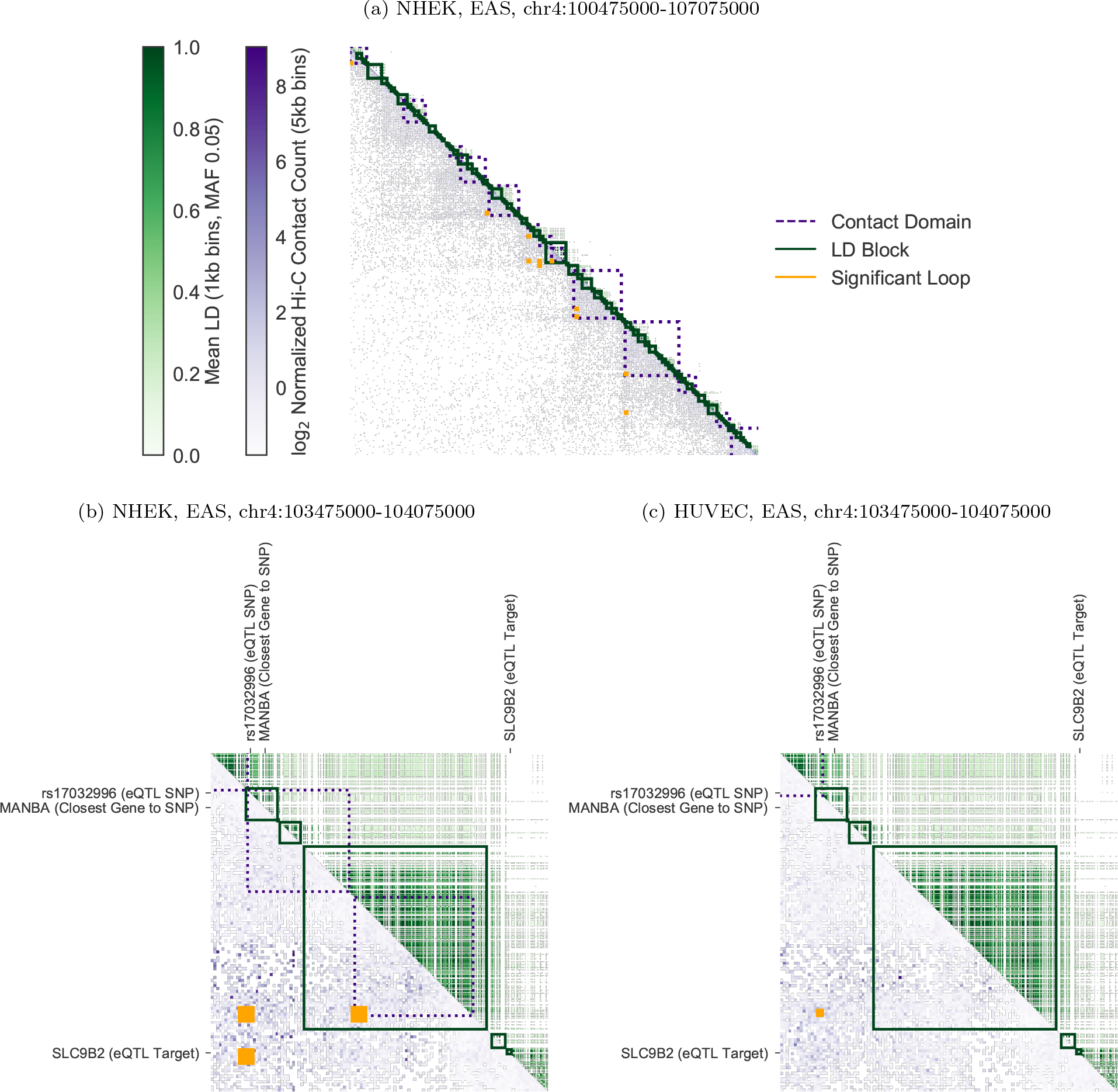
Discordance between LD and Hi-C. An annotated matrix illustrates differences between the genomic scales of LD [24] (*R*^2^, upper triangle, green) versus Hi-C contact frequency [21] (lower triangle, purple). Rows and columns are binned genomic coordinates (hg19) with lower bins near the upper left; for example, row 10 column 11 stores the LD between a bin and its neighbor, while row 11 column 10 stores the contact frequency. More frequent contacts (5kb bins) are darker purple; higher LD (averaged over non-zero LD pairs in 1kb bins) are darker green. Contact domains (nested purple squares) and significant interactions (orange squares) were computed from Hi-C data. LD blocks (green squares) were computed from 1000 Genomes genotypes. While some LD blocks fall within contact domains, there are also many cases where they overlap domain boundaries. (a) A representative 6.6mb locus on chromosome 4 shows Hi-C contacts (NHEK cells) span much longer distances than LD (EAS superpopulation). (b) A 600kb locus on the same chromosome illustrates the complexities of mapping a non-coding SNP (rs17032996) to a target gene. The closest gene MANBA falls within the same LD block as the SNP. However, Hi-C data shows the SNP contacts the SLC9B2 gene ≈ 460kb away in NHEK cells, skipping over intervening expressed gene MANBA. rs17032996 is also an eQTL in B cells [25] and significantly interacts with SLC9B2 in several blood cell types [12]. (c) In HUVEC cells, the SNP no longer interacts with SLC9B2 and several contact domains are lost.

**Table 1:** Accuracy of methods for mapping non-coding regions to genes. Using PIRs from statistically significant PCHi-C interactions in 17 blood cell types as a gold standard [12], we computed the percent agreement of various heuristics for mapping non-coding regions to all baited promoters genome-wide. The closest gene method predicts that a non-coding region interacts with the nearest promoter. The LD heuristic predicts that a non-coding region interacts with any gene in its LD block (one prediction per 1000 Genomes superpopulation). The agreement of LD is even lower than that of the closest gene: 2-7% versus 7-12% across superpopulations.

| Cell Line | Closest Gene | LD (AFR) | LD (AMR) | LD (EAS) | LD (EUR) | LD (SAS) |
| --- | --- | --- | --- | --- | --- | --- |
| Mon | 9.9% | 2.1% | 4.3% | 5.5% | 5.0% | 4.8% |
| Mac0 | 9.4% | 1.7% | 3.5% | 4.4% | 4.2% | 4.0% |
| Mac1 | 10.1% | 2.1% | 4.4% | 5.4% | 5.1% | 4.9% |
| Mac2 | 10.4% | 2.2% | 4.6% | 5.6% | 5.5% | 5.2% |
| Neu | 9.8% | 2.5% | 5.1% | 6.3% | 6.0% | 5.4% |
| MK | 11.9% | 3.0% | 5.9% | 7.4% | 6.9% | 6.6% |
| EP | 10.9% | 2.4% | 4.8% | 6.0% | 5.7% | 5.4% |
| Ery | 8.7% | 2.8% | 5.7% | 7.0% | 6.7% | 6.3% |
| FoeT | 7.0% | 2.0% | 4.1% | 5.1% | 4.8% | 4.6% |
| nCD4 | 8.8% | 2.4% | 5.0% | 6.3% | 5.9% | 5.6% |
| tCD4 | 7.9% | 2.0% | 4.1% | 5.2% | 4.9% | 4.7% |
| aCD4 | 9.3% | 2.3% | 4.9% | 6.2% | 5.8% | 5.5% |
| naCD4 | 8.6% | 2.0% | 4.3% | 5.4% | 5.1% | 4.8% |
| nCD8 | 8.4% | 2.4% | 4.8% | 6.1% | 5.7% | 5.4% |
| tCD8 | 9.2% | 2.7% | 5.3% | 6.7% | 6.2% | 5.9% |
| nB | 9.0% | 2.2% | 4.4% | 5.8% | 5.4% | 5.1% |
| tB | 8.7% | 2.2% | 4.2% | 5.6% | 5.2% | 4.9% |

### LD is low between distal regulatory SNPs and their genes

Genetic variants associated with statistically significant differences in a gene’s expression (eQTLs) provide evidence of functional relationships between regulatory regions and genes separated by long genomic distances. Indeed, target genes of GTEx eQTLs [29] and blood eQTLs [25] have median distances of 49 and 113kb, respectively. Combined with our other findings, these distances suggest that a distal eQTL and its target gene are likely to have zero LD and thus be separated by a large number of LD blocks. We therefore compared the frequency of eQTLs amongst non-coding regions that interact with gene promoters versus distance-matched regions that do not. Analyzing B-cells where both PCHi-C [12] and eQTL [25] data were available, statistically significant chromatin interactions were highly enriched for eQTLs across genomic distances up to 1.6mb (Figure 6, Figure 1). In contrast, regions in strong LD with a promoter were only enriched for eQTLs at genomic distances less than 100kb, emphasizing that more distal eQTLs are often in 3D proximity to their target promoters but are not genetically linked to them. This result holds across all super-populations. Thus, genomic distance and LD are poor predictors of the genes targeted by distal eQTLs [11].

### Mapping non-coding variants to genes with Hi-C produces more functional enrichments than genomic distance or LD

If regulatory interactions are common at large genomic distances where LD is approximately zero, then GWAS SNPs linked to genes via Hi-C should include more true gene targets than using closest genes or genes in LD with the SNP. If true, then the set of genes associated with GWAS hits via Hi-C should also share more functional annotations. To test this idea, we examined the magnitude and statistical significance of Gene Ontology (GO) enrichments for genes associated with all GWAS SNPs for a given phenotype via PCHi-C interactions [12] (all genes with promoter interacting regions overlapping the SNP), genomic distance (closest promoter to the SNP), or genetic distance (all promoters in the same LD-block as the SNP). Most blood-relevant phenotypes in the GWAS catalog [30] had the largest number of significantly enriched GO terms using blood cell PCHi-C assignment (Figure 7), even after controlling for the number of genes mapped to GWAS SNPs. LD-based assignment occasionally produced a limited number of terms with effect sizes larger than those from PCHi-C assignment (Figure 7c). Nonetheless, using LD resulted in fewer GO terms associated with the phenotype and a lower area-under-the-curve than PCHi-C. As a negative control, we examined GWAS SNPs for phenotypes not relevant to blood and found that closest gene and LD approaches have more GO enrichments than for blood-relevant phenotypes, confirming that PCHi-C enrichments are tissue-specific. Thus, PCHi-C data from blood does not correctly associate SNPs with genes for non-blood phenotypes, highlighting the need for interaction data collected in the cell type of interest to harness the power of chromatin interactions for functional assignment.

**Figure 3:**
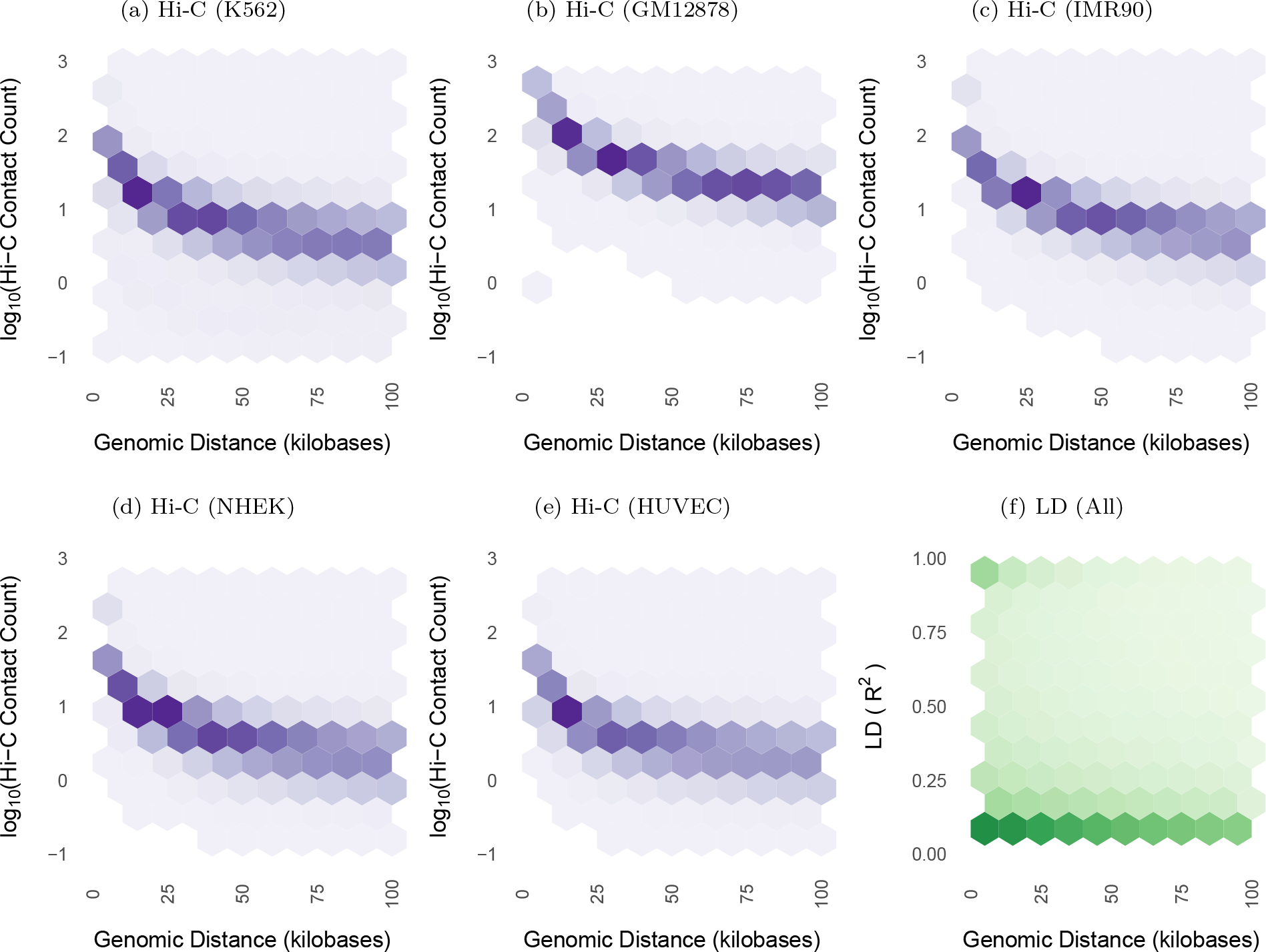
LD and Hi-C contact decay with genomic distance. Both Hi-C contact frequency [21] (panels a-e) and LD (panel f) are anti-correlated with genomic distance (Spearman *ρ* between −0.5 and −0.71 for Hi-C across cell lines; *ρ* ≈ −0.52 for LD). All plots display non-zero values from their respective datasets. LD decays towards zero at much shorter genomic distance than contact frequency, with high LD SNP pairs concentrated below 50kb. In contrast, Hi-C contacts are common up to and exceeding the median length of contact domains (250kb) or TADs (840kb). Supplementary Figure 2 highlights decay up to 2mb, while this figure highlights decay up to 100kb. Supplementary Figure 4 shows nearly identical LD scaling per superpopulation. Contact frequencies vary in approximate proportion to sequencing depth and number of replicates per cell line (Supplementary Table 1). Panel f) shows 836 million biallelic SNPs on chromosome 14 and is representative of other chromosomes.

**Figure 4:**
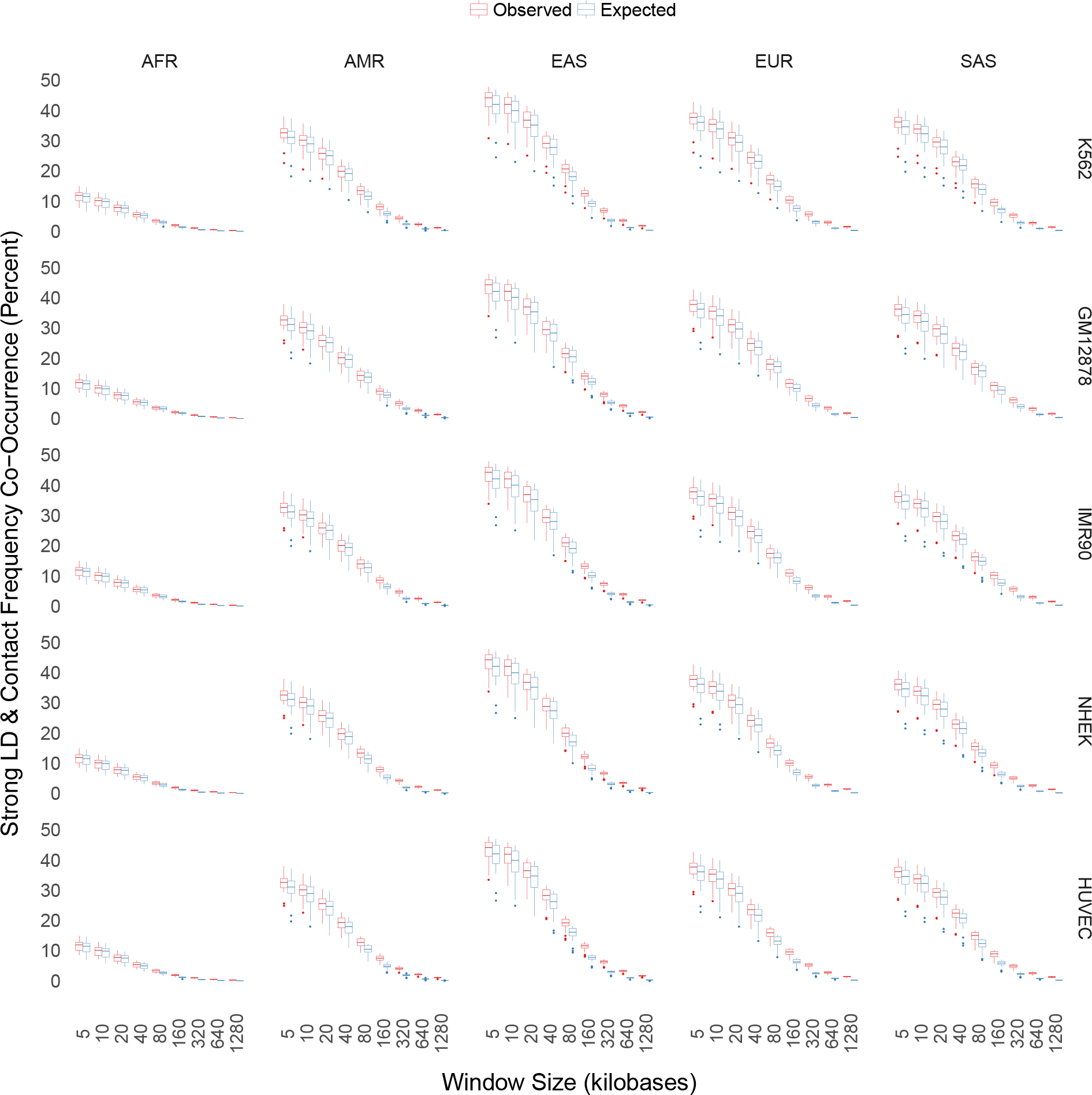
Genome-wide Hi-C and LD concordance. Frequent Hi-C contacts and strong LD (*R*^2^ > 0.8) co-occur less than 50% of the time at short genomic distances. Concordance is cut nearly in half at 40kb where most LD has decayed to 0, and is nearly 0 at many scales where statistically significant chromatin interations occur. In addition, concordance and rate of decay varies by super-population, with AFR having only ≈ 12% concordance at short genomic distances

**Figure 5:**
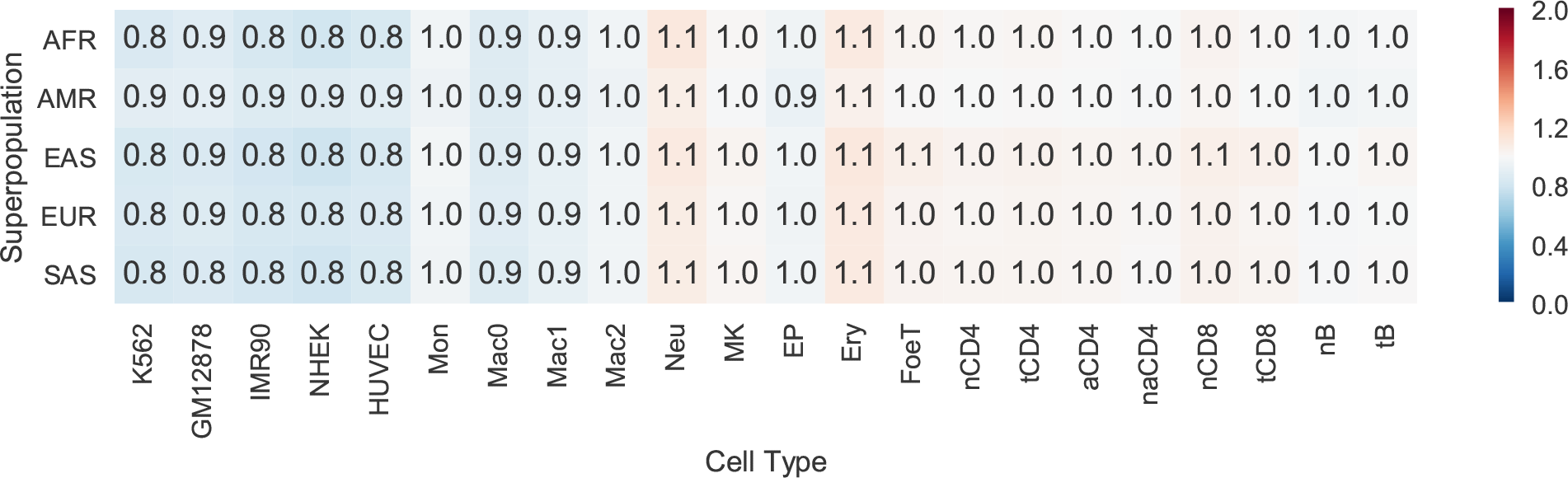
Significant versus non-significant chromatin interaction LD. For SNPs located on the fragments of statistically significant and distance-matched non-significant chromatin interactions, the maximum pairwise LD between SNPs (*interaction LD*) was computed for 5 Hi-C and 17 PCHi-C datasets. The ratio of mean interaction LD for significant versus non-significant interactions quantifies how well LD acts as a proxy for chromatin interactions; a ratio greater than 1 indicates significant interactions are enriched for SNPs in strong LD. However, this ratio is near 1 for all cell types and superpopulations, indicating that LD is not a sufficient proxy for chromatin interactions. Supplemental Table 2 provides raw values for this figure; ratios smaller or larger than 1 are the result of relatively small differences in weak LD.

**Figure 6:**
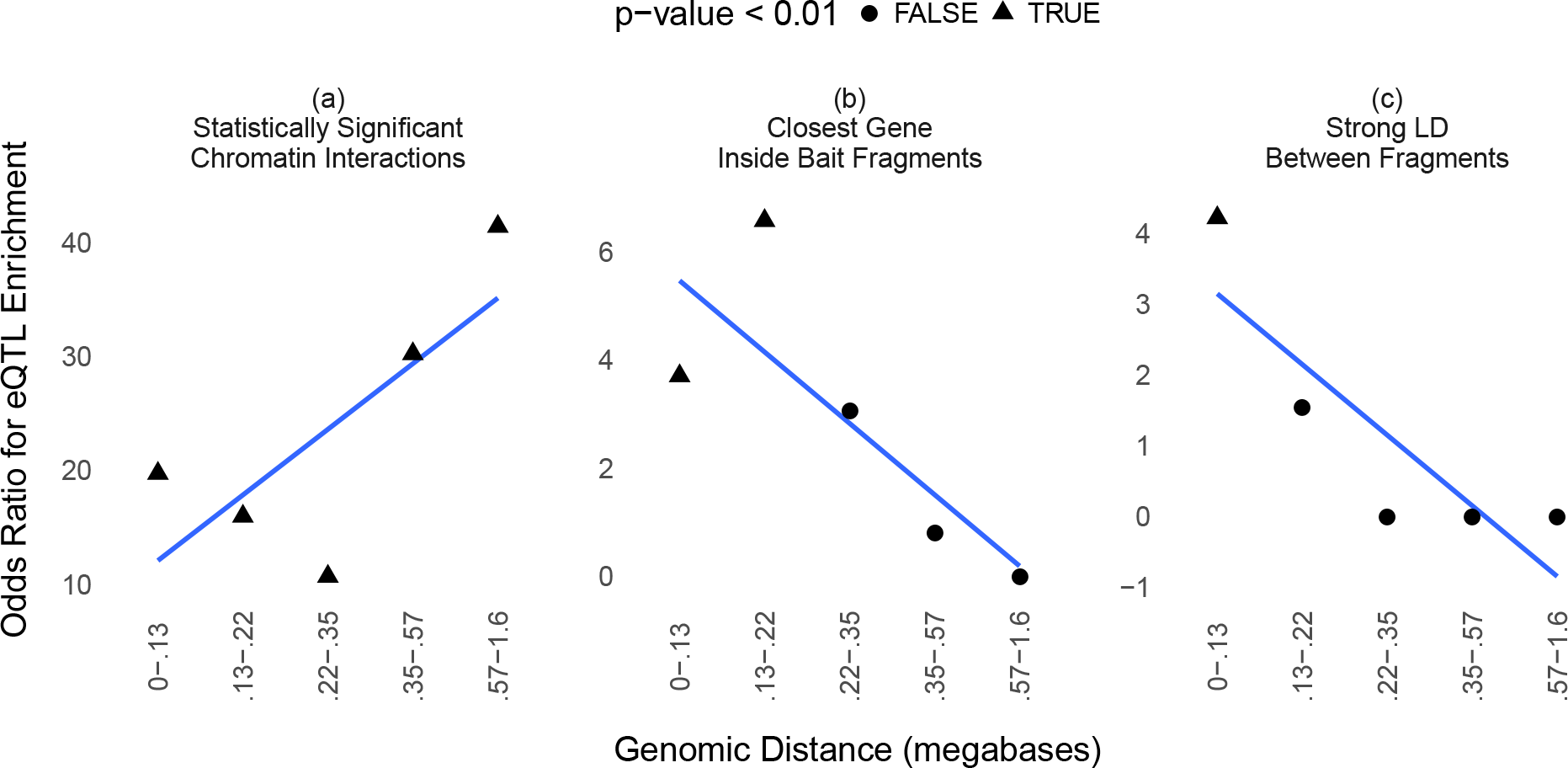
eQTL enrichment. B-cell eQTLs [25] are enriched across quantile-binned distances in statistically significant B-cell PCHi-C interactions [12] (panel a). Promoter-interacting fragments with bait fragments containing the closest gene are enriched for eQTLs at shorter distance bins (panel b), while interacting fragments in strong LD (maximum pairwise *R*^2^ > 0.8, averaged across superpopulations) are only enriched at the most proximal bin (panel c).

**Figure 7:**
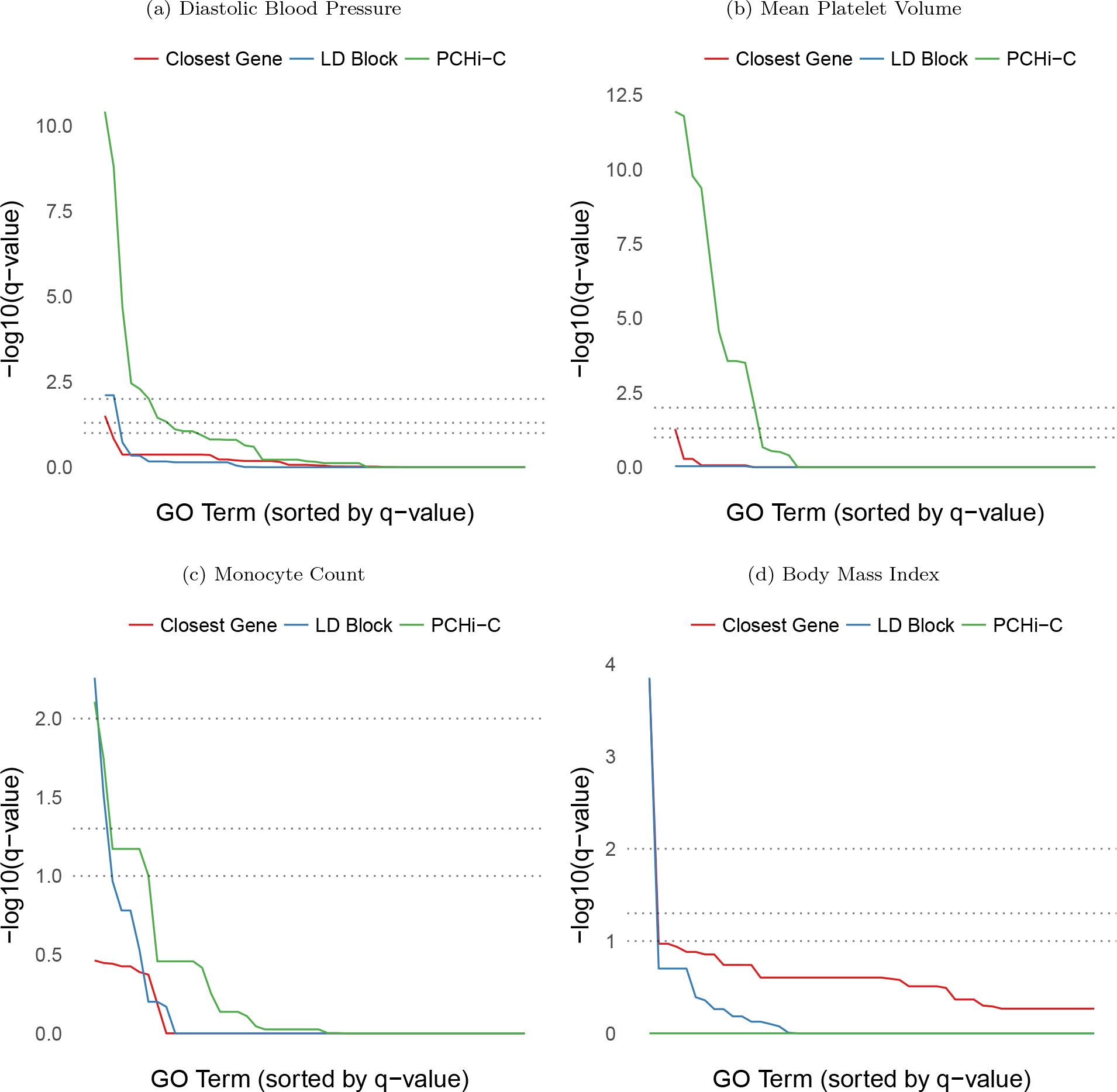
GO enrichment for closest, same-LD-block, and PCHi-C bait genes. Enrichment of GO terms (Benjamini-Hochberg adjusted p-values, −log_10_ scale) in multiple blood-relevant phenotypes from the GWAS catalog [30]. Methods for functional assignment of SNPs include using the closest gene, all genes within the same LD block (EUR super-population), and promoter capture bait genes with a SNP located in the promoter-interacting region of a statistically significant blood cell chromatin interaction [12]. Gray horizontal lines indicate FDR cutoffs of 1, 5, and 10 percent. In blood-relevant phenotypes (panels a-c), PCHi-C bait genes interacting in 3D with GWAS SNPs typically show more enrichment for a larger number of terms than same-LD-block or closest gene approaches, whose enrichment is affected by large numbers of false positives and negatives. For non-blood phenotypes, chromatin interactions in the wrong cell type can have little or no enrichment compared to closest gene and same-LD-block approaches (panel d).

## Discussion

Chromatin interactions and LD are both pairwise measurements between genomic loci that show block patterns along mammalian chromosomes. Given their similar structure, it might be tempting to speculate that LD blocks correspond to or are contained within three-dimensional chromatin domains. In this study, we conducted diverse genome-wide analyses across different length scales and showed that LD maps are not correlated with chromatin interaction maps. The main factor driving these differences is the frequency of chromatin interactions over genomic distances where the genetic linkage between SNPs is close to zero. Significant chromatin interactions often span hundreds or sometimes thousands of LD blocks. However, even at genomic distances where contact frequencies and LD are both high on average, the correlation between their block patterns is weak. This discordance is perhaps not surprising given the different origins of chromatin domains (insulated regulatory regions) and LD blocks (recombination), as well as the increasing number of examples of distal enhancers and eQTLs documented in the literature. Despite growing awareness that regulatory interactions need not be in LD or nearby on the genome, this phenomenon appears not to have been quantified systematically genome-wide until now. To do so, we developed computationally efficient statistical analyses and leveraged 5 ENCODE cell lines [21], 17 human primary blood cell types [12], and 5 superpopulations from the 1000 Genomes Project [24]. The results document that LD and chromatin interactions are indeed uncorrelated at genomic distances of 5kb or more.

This study has important implications for associating non-coding variants with genes, downstream phenotypes, and molecular mechanisms. Our results show that variants have great potential to regulate genes beyond their LD block. Hence, the common practice of mapping candidate regulatory variants to the closest gene or other genes in the same LD block will typically miss most chromatin interactions between the variant and gene promoters. In addition, LD is the same in all cell types whereas within-TAD regulatory interactions vary across cell types [21] (Figure 2, Supplementary Figure 1). Despite the emergence of chromatin interaction data as a new paradigm for mapping non-coding variants to genes, LD blocks and genomic distance are still widely used, resulting in many GWAS hits being annotated with incorrect genes. There are good reasons why genomic distance and LD have been used to annotate regulatory variants: these quantities are relatively easy to compute, and only a few cell types currently have high enough resolution chromatin interaction data (≈ 1-5kb) for linking specific regulatory variants to promoters. The lack of correlation between these common heuristics and high resolution chromatin contacts underscores the importance of generating or computationally predicting chromatin structure across many more cell types.

In addition to highlighting the need for incorporating chromatin interactions into functional assignment, the discordance between chromatin contact frequency and LD has evolutionary implications. One consequence is that entire TADs or sub-TADs do not typically segregate as single haplotypes in human populations, enabling independent selection on regulatory variants versus the promoter and coding variants of their target genes. Furthermore, recombination blocks that span chromatin domain boundaries indicate that regulatory and coding variants from one domain can segregate with variants from the adjacent domain. The fact that haplotype breakpoints do not align with chromatin boundaries may indicate that recombination is deleterious at these functional elements, perhaps due to the mutagenic effects of recombination. These findings are different from observations regarding fixed structural differences between genomes of different mammals, which tend to preserve TADs with breakpoints enriched at TAD boundaries [31, 32]. We therefore conclude that while chromatin domains are functional genomic entities maintained as syntenic units over evolutionary time, recombination is independent of chromatin structure. This creates novel haplotypes of the genomic elements within and between TADs on which selection can operate.

## Methods

In order to perform large scale analyses, some caveats were necessary in order to place reasonable bounds on compute time and memory, even in a high-performance computing environment. For example, LD was computed between SNP pairs at most 2mb apart and stored if LD was 0.01 or greater. Also, the resolution of Hi-C and PCHi-C data prevented examining correlations between chromatin interactions and LD at genomic distances below 5kb. To make our choices transparent and our analyses reproducible, our code is available at https://github.com/shwhalen/loopdis.git.

Hi-C data [21] was obtained from the NCBI Gene Expression Omnibus using accession GSE63525 (including contact domains, statistically significant loops, and sparse contact matrices along with coefficients for normalization and expectation). Promoter capture Hi-C data [12] was obtained from Open Science Framework (https://osf.io/u8tzp/). All data sources use human reference genome hg19. Analyses utilized bcftools 1.6, bedtools 2.27.1 [33], plink 1.90b5 [34], pandas 0.22.0 [35], matplotlib 2.1.1 [36], seaborn 0.8.1, ggplot 2.2.1 [37], and GNU Parallel 20171222 [38]. Python 3.6.4 was provided by the Miniconda distribution; R 3.4.3 was compiled from source using gcc 7.2.1.

### Linkage Disequilibrium

Bi-allelic SNPs from phase 3 of the 1000 Genome Project were first converted to plink’s binary BED format (`--make-bed --allow-extra-chr --biallelic-only`), and the pairwise LD computed (`--r2`) for all SNPs with a minimum MAF of 5% (`--maf 0.05`) located within 2mb of each other (`--ld-window-kb 2000`). The number of pairwise comparisons allowed within a window was increased (`--ld-window 10000`), and the default *R*^2^ filter lowered from 20% down to 1% (`--ld-window-r2 0.01`). Pairs below this threshold were assigned *R*^2^ = 0. LD computations were performed separately for each superpopulation by using the 1000 Genomes panel file (`integrated call samples v3.20130502.ALL.panel`) and the `--filter` flag.

LD blocks were computed using plink (utilizing the algorithm from Gabriel et al. [28]) with the `--blocks no-pheno-req no-small-max-span --blocks-max-kb 2000` flags. Blocks were computed separately for each superpopulation using the same `--filter` flag and panel file.

### Interacting versus Non-Interacting LD

For each 1000 Genomes superpopulation, bi-allelic SNPs with a minimum MAF of 5% were intersected with either Hi-C [21] or promoter capture Hi-C [12] fragments using the `bedtools pairtobed` command with the `-type both` flag. For each pair of interacting fragments, the maximum LD between SNPs on different fragments was computed. The mean of this maximum pairwise LD was computed separately for statistically significant and non-significant interactions in order to compute a ratio.

For Hi-C data, negatives (non-significantly interacting fragments) were obtained by shuffling a list of positives (significant interactions) called by the Juicer pipeline at 10% FDR [26] along the same chromosome. For promoter capture Hi-C data, positives and negatives were obtained from a list of interactions scored by the CHiCAGO pipeline [27]. As in the original paper, negatives were interactions with a score less than 5. For both datasets, negatives were distance-matched to positives using quantile binning of interaction distance.

### Hi-C versus LD Concordance

Observed over expected Hi-C values were computed using formulas from Rao et al. [21] applied to VC-normalized contact counts at 5kb resolution for each ENCODE cell line. For comparable resolution, LD per 5kb genomic bin was computed for each 1000 Genomes superpopulation using the 75th percentile of pairwise LD values in the bin. This was more robust outliers and heavily zero-skewed LD distributions than the average or median.

Concordance was computed based on whether a bin’s LD value was strong (*R*^2^ > 0.8) and its chromatin contact frequency was strong (above the 75th percentile of contact frequencies), for all bins located in non-overlapping genomic windows of fixed size. This was repeated for window sizes of 5, 10, 20, 40, 80, 160, 320, 640, and 1280kb to examine concordance across multiple scales, and without variation introduced by different TAD-calling algorithms.

### eQTL Statistics

B-cell eQTLs [25] were intersected with naive B-cell promoter capture Hi-C interactions [12]; the eQTL was required to overlap the promoter-interacting region and the eQTL target was required to overlap the bait fragment. The presence or absence of an interacting eQTL was stored in a binary vector. Next, the closest gene to each promoter-interacting fragment was computed using `bedtools closest` and Ensembl gene annotations. The presence or absence of the closest gene in the corresponding bait fragment was stored in a binary vector. Next, the statistical significance of chromatin interactions (thresholded using a CHiCAGO score of 5) was stored in a binary vector. Finally, for each superpopulation, a binary vector stored the maximum pairwise LD between fragments.

eQTLs were tested for enrichment in statistically significant chromatin interactions, interactions where the bait was the closest gene, and interactions where the maximum pairwise LD between fragments was > 0.8 (averaged over superpopulations). Interactions were quantile binned by distance up to 1.6mb so that all entries in the contingency table would be non-zero. For each distance bin, the odds ratio was computed as the ratio of the diagonal to the off-diagonal entries of the contingency table, and the p-value was computed using R’s `fisher.test` function.

### Gene Ontology Enrichment

The promoter-interacting region of statistically significant PCHi-C interactions [12] was intersected with SNPs for the 30 most abundant phenotypes in the GWAS catalog [30] (release 2018-01-31). For each GO term, a Fisher’s exact test was computed on a 2 by 2 contingency table counting if the interaction contained a GWAS SNP for the phenotype in its PIR, and whether or not the interaction’s bait gene was annotated with that GO term. Benjamini-Hochberg correction for multiple hypothesis testing was applied to the resulting p-values. For comparison, this was repeated for the closest gene to each GWAS SNP, as well as all genes in the same LD block as the GWAS SNP.

## Acknowledgments

Thanks to Dr. Hunter Fraser and Dr. Jonathan Pritchard for GO analysis suggestions. Dr. Noah Zaitlen, Dr. Dan Geschwind, Dr. Hyejung Won, and Dr. Marisa Wong Medina provided helpful feedback. This project was supported by the Bench to Bassinet Program of the NHLBI (U01HL098179, UM1HL098179), NIH/NHLBI (HL089707), NIH/NIMH (MH109907), the San Simeon Fund, and the Gladstone Institutes.

## Author contributions

SW and KSP designed the experiments; SW coded the experiments; SW and KSP wrote the paper.

## Supplemental Material

### Supplemental Tables

**Supplementary Table 1:**
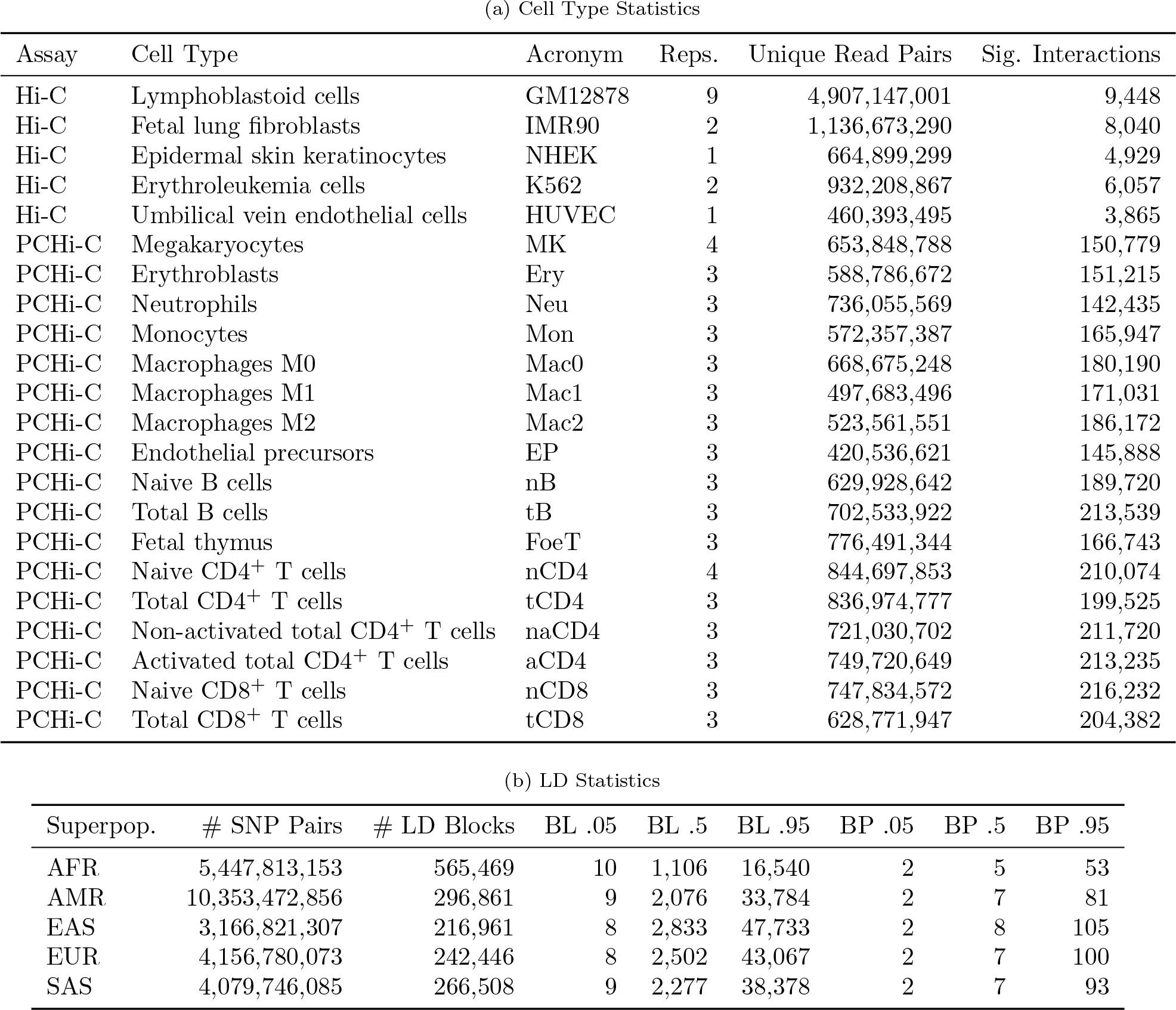
Dataset statistics. a) Summary of Hi-C [21] and promoter capture Hi-C (PCHi-C) [12] datasets. Statistically significant (sig.) Hi-C interactions were called on combined replicates (reps.) by the Juicer pipeline with a 10% false discovery rate (FDR) [26], while PCHi-C interactions were scored by the CHiCAGO pipeline [27] with a significance threshold of 5. PCHi-C has 15-17 fold enrichment for promoter interactions, resulting in an effective coverage of 165 billion read pairs compared to 15 billion for Hi-C [12]. b) Number of unique SNP pairs and LD blocks per superpopulation. LD block quantiles (.05, .5, and .95) for length (BL) and unique SNP pairs (BP) are also given. SNPs were filtered as described in Methods.

| Assay | Cell Type | Acronym | Reps. | Unique Read Pairs | Sig. Interactions |
| --- | --- | --- | --- | --- | --- |
| Hi-C | Lymphoblastoid cells | GM12878 | 9 | 4,907,147,001 | 9,448 |
| Hi-C | Fetal lung fibroblasts | IMR90 | 2 | 1,136,673,290 | 8,040 |
| Hi-C | Epidermal skin keratinocytes | NHEK | 1 | 664,899,299 | 4,929 |
| Hi-C | Erythroleukemia cells | K562 | 2 | 932,208,867 | 6,057 |
| Hi-C | Umbilical vein endothelial cells | HUVEC | 1 | 460,393,495 | 3,865 |
| PCHi-C | Megakaryocytes | MK | 4 | 653,848,788 | 150,779 |
| PCHi-C | Erythroblasts | Ery | 3 | 588,786,672 | 151,215 |
| PCHi-C | Neutrophils | Neu | 3 | 736,055,569 | 142,435 |
| PCHi-C | Monocytes | Mon | 3 | 572,357,387 | 165,947 |
| PCHi-C | Macrophages M0 | Mac0 | 3 | 668,675,248 | 180,190 |
| PCHi-C | Macrophages M1 | Mac1 | 3 | 497,683,496 | 171,031 |
| PCHi-C | Macrophages M2 | Mac2 | 3 | 523,561,551 | 186,172 |
| PCHi-C | Endothelial precursors | EP | 3 | 420,536,621 | 145,888 |
| PCHi-C | Naive B cells | nB | 3 | 629,928,642 | 189,720 |
| PCHi-C | Total B cells | tB | 3 | 702,533,922 | 213,539 |
| PCHi-C | Fetal thymus | FoeT | 3 | 776,491,344 | 166,743 |
| PCHi-C | Naive CD4<sup>+</sup> T cells | nCD4 | 4 | 844,697,853 | 210,074 |
| PCHi-C | Total CD4<sup>+</sup> T cells | tCD4 | 3 | 836,974,777 | 199,525 |
| PCHi-C | Non-activated total CD4<sup>+</sup> T cells | naCD4 | 3 | 721,030,702 | 211,720 |
| PCHi-C | Activated total CD4<sup>+</sup> T cells | aCD4 | 3 | 749,720,649 | 213,235 |
| PCHi-C | Naive CD8<sup>+</sup> T cells | nCD8 | 3 | 747,834,572 | 216,232 |
| PCHi-C | Total CD8<sup>+</sup> T cells | tCD8 | 3 | 628,771,947 | 204,382 |

Supplementary Table 1: **Dataset statistics** a) Summary of Hi-C [21] and promoter capture Hi-C (PCHi-C) [12] datasets.
| Superpop. | # SNP Pairs | # LD Blocks | BL .05 | BL .5 | BL .95 | BP .05 | BP .5 | BP .95 |
| --- | --- | --- | --- | --- | --- | --- | --- | --- |
| AFR | 5,447,813,153 | 565,469 | 10 | 1,106 | 16,540 | 2 | 5 | 53 |
| AMR | 10,353,472,856 | 296,861 | 9 | 2,076 | 33,784 | 2 | 7 | 81 |
| EAS | 3,166,821,307 | 216,961 | 8 | 2,833 | 47,733 | 2 | 8 | 105 |
| EUR | 4,156,780,073 | 242,446 | 8 | 2,502 | 43,067 | 2 | 7 | 100 |
| SAS | 4,079,746,085 | 266,508 | 9 | 2,277 | 38,378 | 2 | 7 | 93 |

**Supplementary Table 2:**
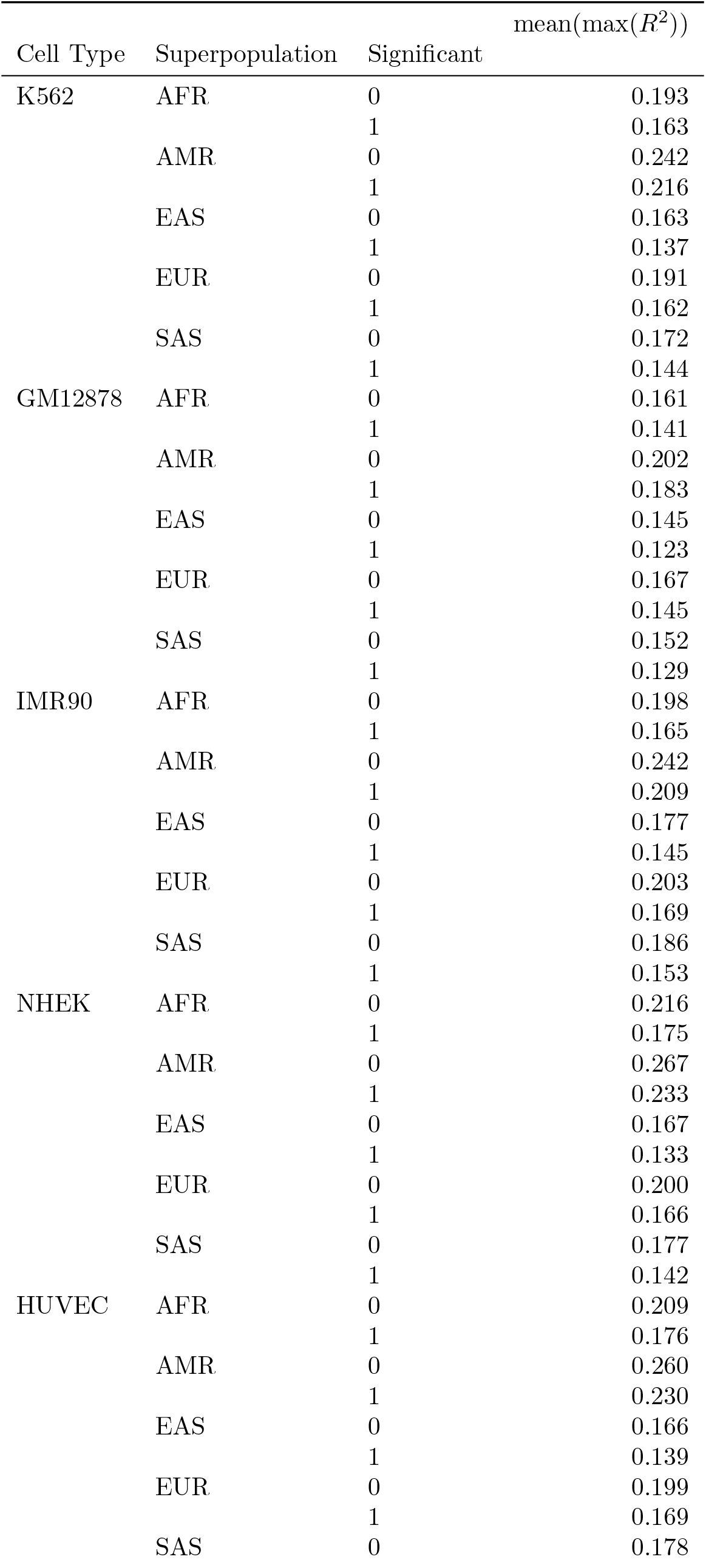
Significant versus non-significant interaction LD. For SNPs located on the fragments of statistically significant and distance-matched non-significant chromatin interactions, the maximum pairwise LD between SNPs (*interaction LD*) was computed for 5 Hi-C and 17 PCHi-C datasets. The mean interaction LD per cell type is given for statistically significant (1) and non-significant (0) interactions. These means were used to compute the ratios shown in Figure 5.

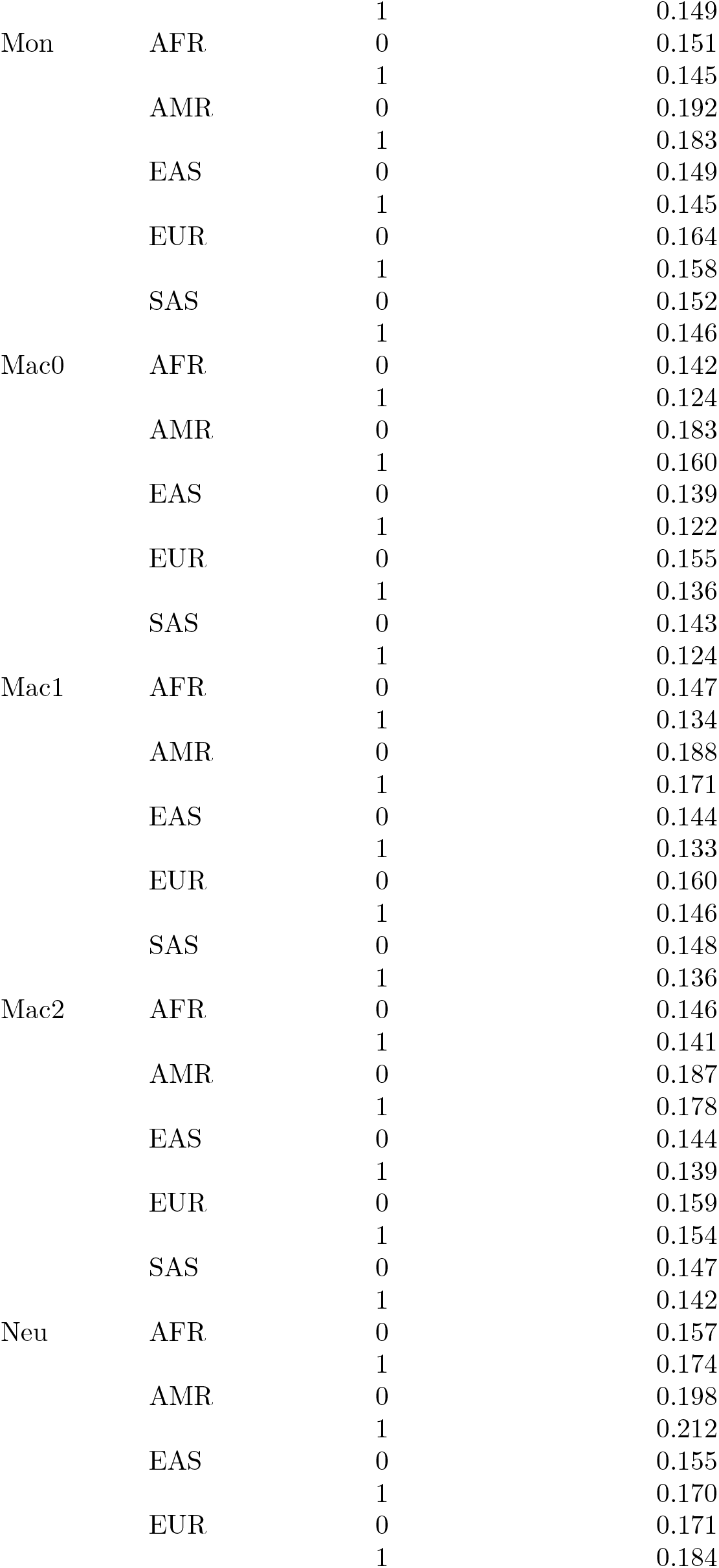

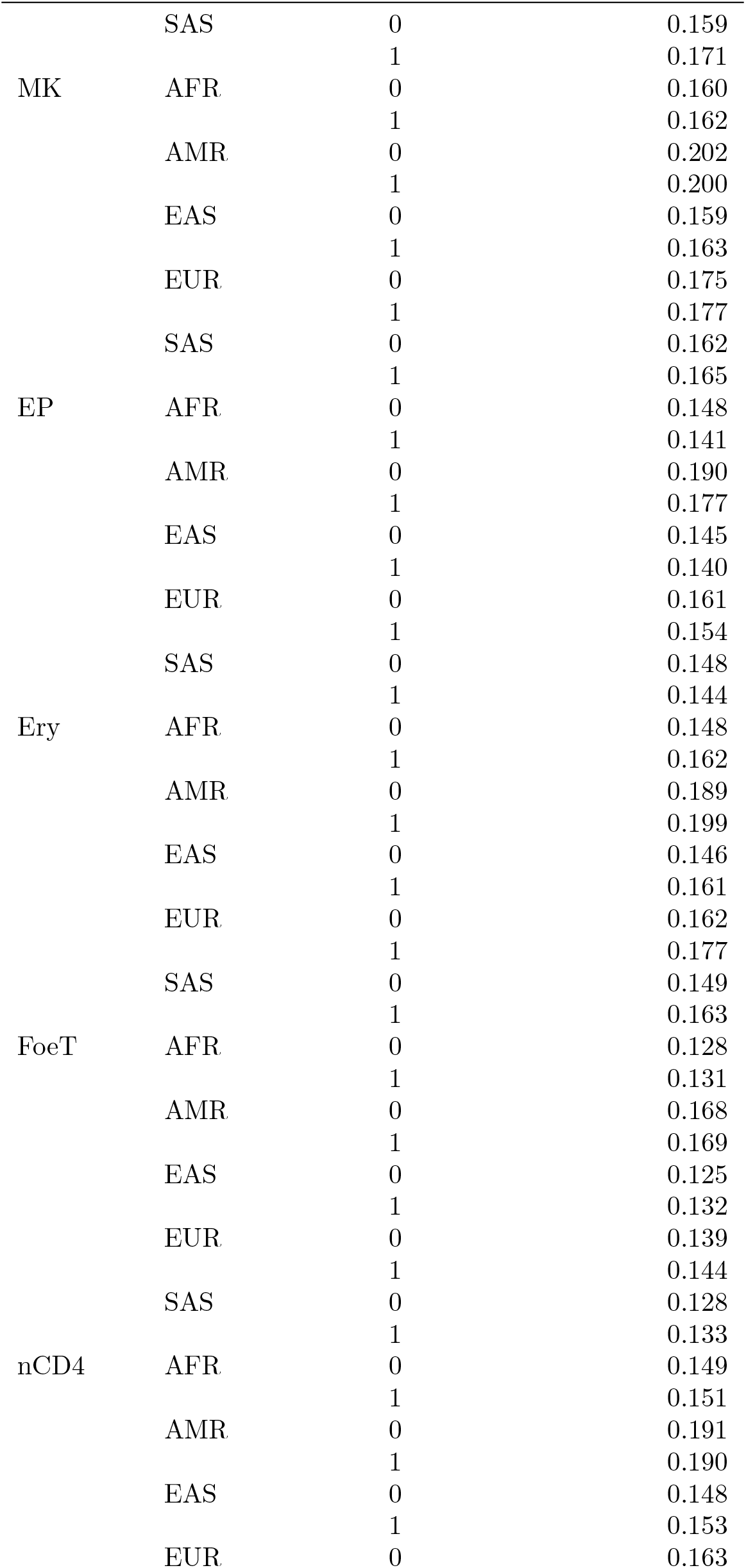

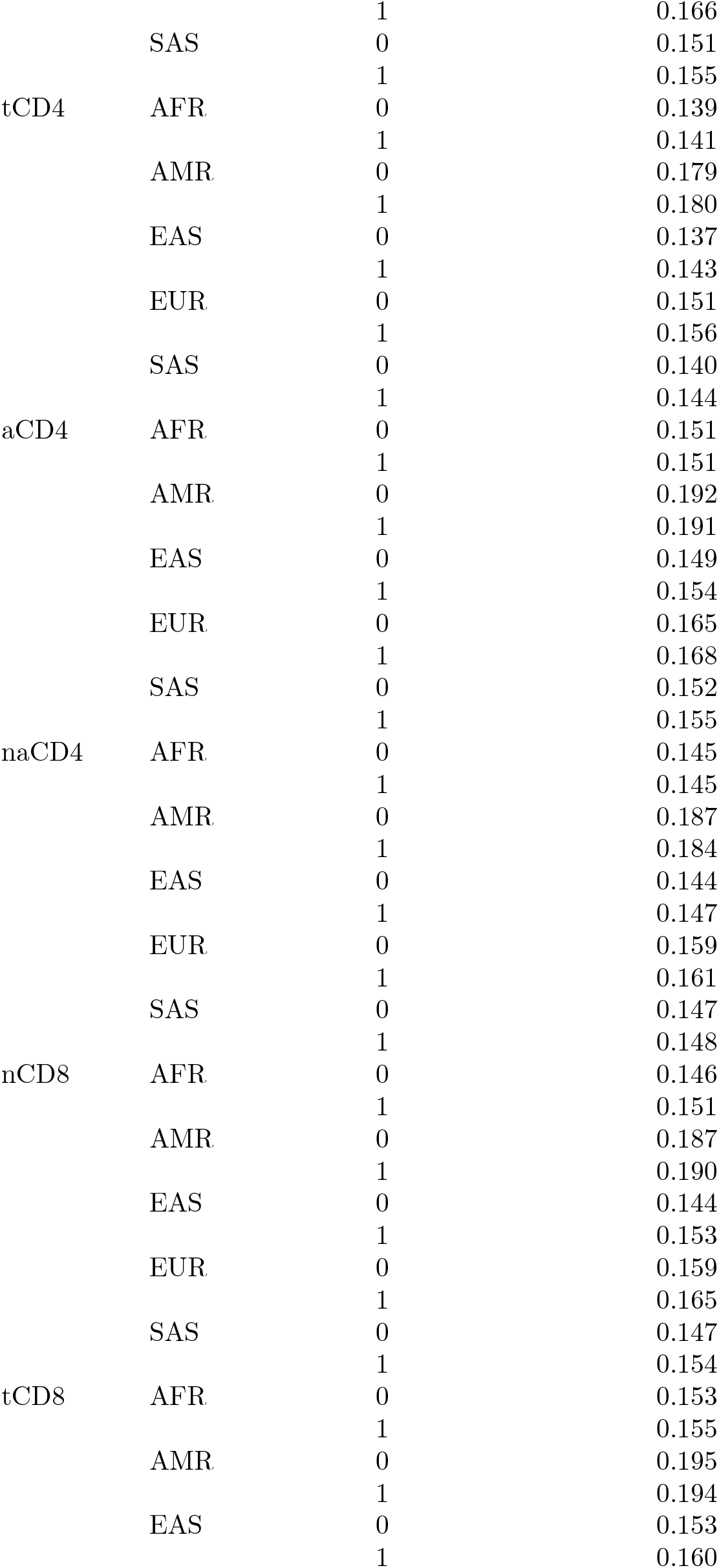

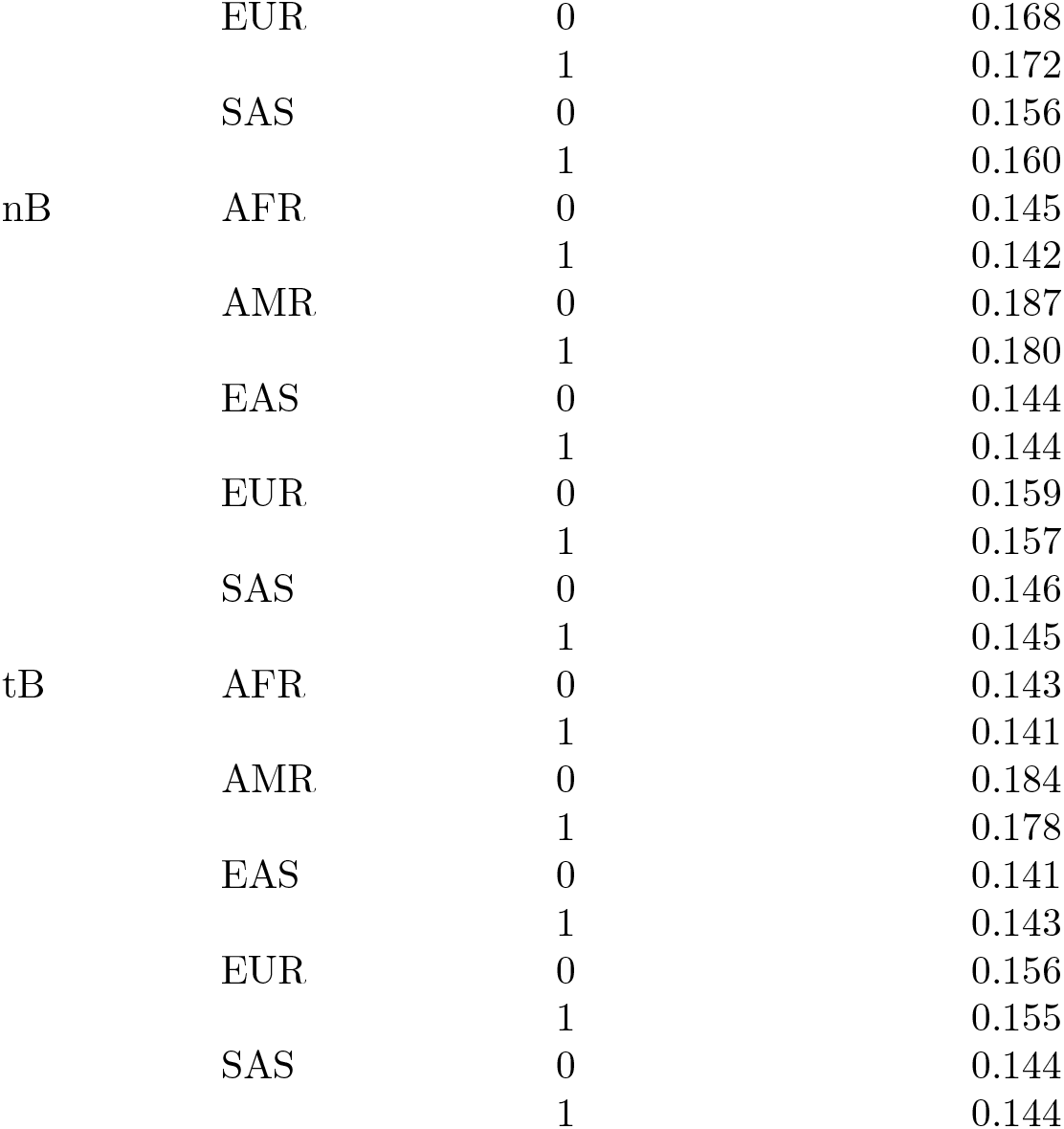

### Supplemental Figures

**Supplementary Figure 1:**
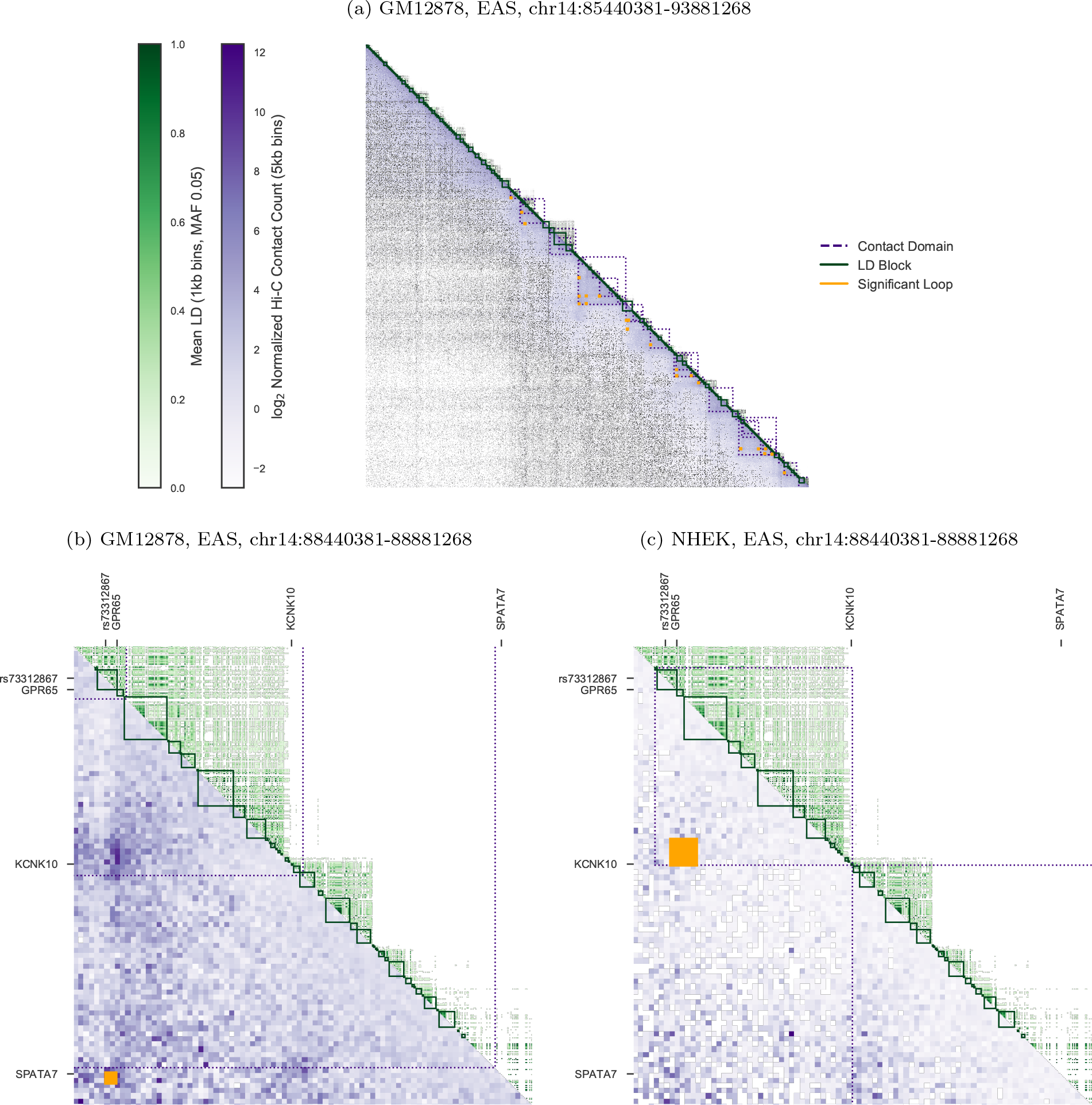
Discordance between LD and Hi-C. An annotated matrix illustrates differences between the genomic scales of LD [24] (*R*^2^, upper triangle, green) versus Hi-C contact frequency [21] (lower triangle, purple). Rows and columns are binned genomic coordinates (hg19) with lower bins near the upper left; for example, row 10 column 11 stores the LD between a bin and its neighbor, while row 11 column 10 stores the contact frequency. More frequent contacts (5kb bins) are darker purple; higher LD (averaged over non-zero LD pairs in 1kb bins) are darker green. Contact domains (nested purple squares) and significant interactions (orange squares) were computed from Hi-C data. (a) A representative 8.5mb locus on chromosome 14 shows Hi-C contacts (GM12878 cells) span much longer distances than LD (EAS superpopulation). (b) A 400kb locus on the same chromosome illustrates the complexities of mapping a non-coding SNP (rs73312867) to a target gene. The closest gene GPR65 falls within the same LD block as the SNP. However, Hi-C data shows the SNP contacts the SPATA7 gene ≈ 380kb away, skipping over intervening gene KCNK10. (c) In NHEK cells, the SNP interacts with KCNK10 instead.

**Supplementary Figure 2:**
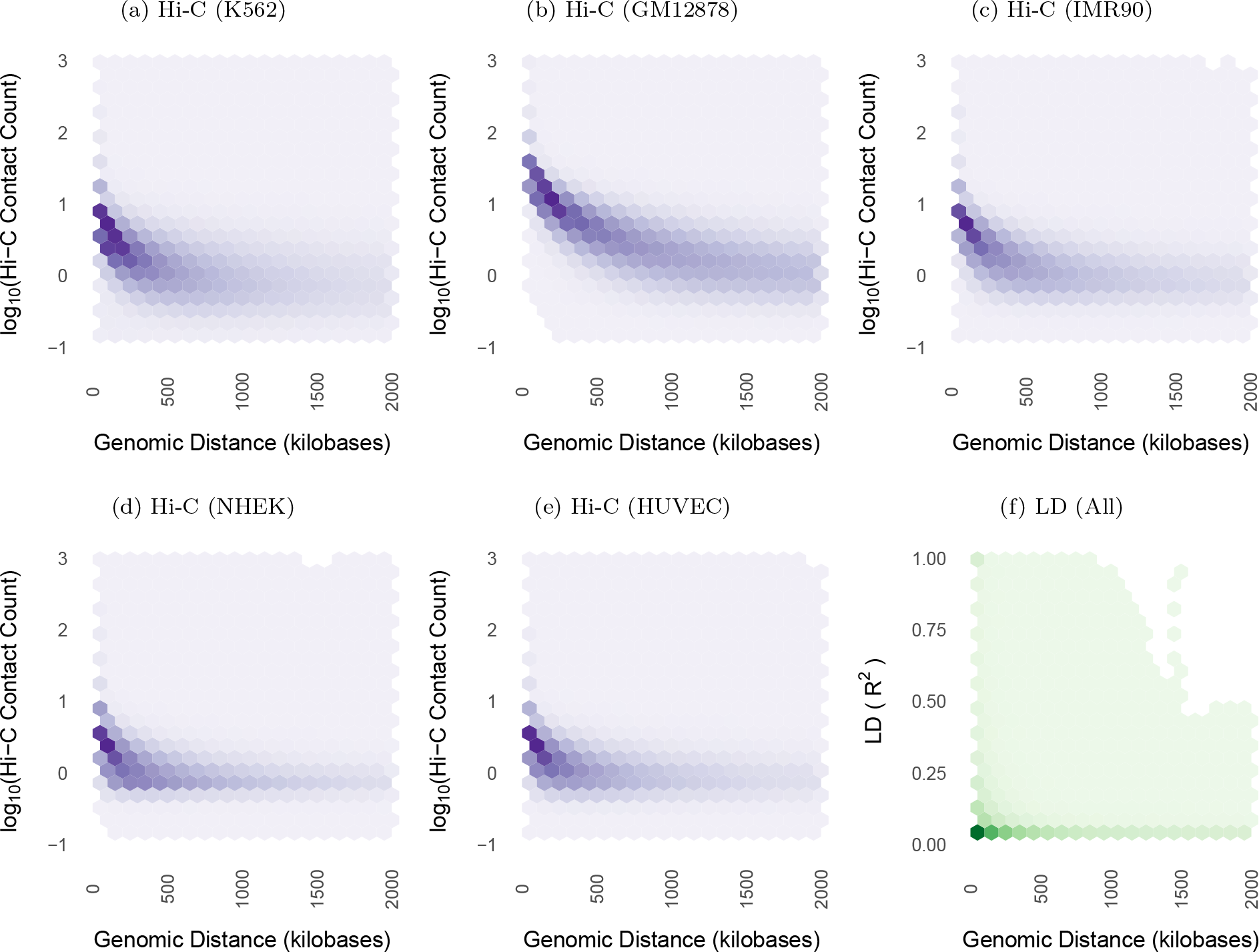
LD and Hi-C contacts decay with genomic distance. Both Hi-C contact frequency [21] (panels a-e) and LD (panel f) are anti-correlated with genomic distance (Spearman *ρ* between −0.5 and −0.71 for Hi-C across cell lines; *ρ* ≈ −0.52 for LD). All plots display non-zero values from their respective datasets. LD decays towards zero at much shorter genomic distance than contact frequency, with most high LD SNP pairs concentrated below 50kb. Hi-C contacts are common at longer genomic distances up to and exceeding the median length of contact domains (250kb) or TADs (840kb). Figure 3 highlights decay up to 100kb, while this figure highlights decay up to 2mb. Supplementary Figure 4 shows nearly identical LD scaling per superpopulation. Panel f) shows 836 million biallelic SNPs on chromosome 14 and is representative of other chromosomes.

**Supplementary Figure 3:**
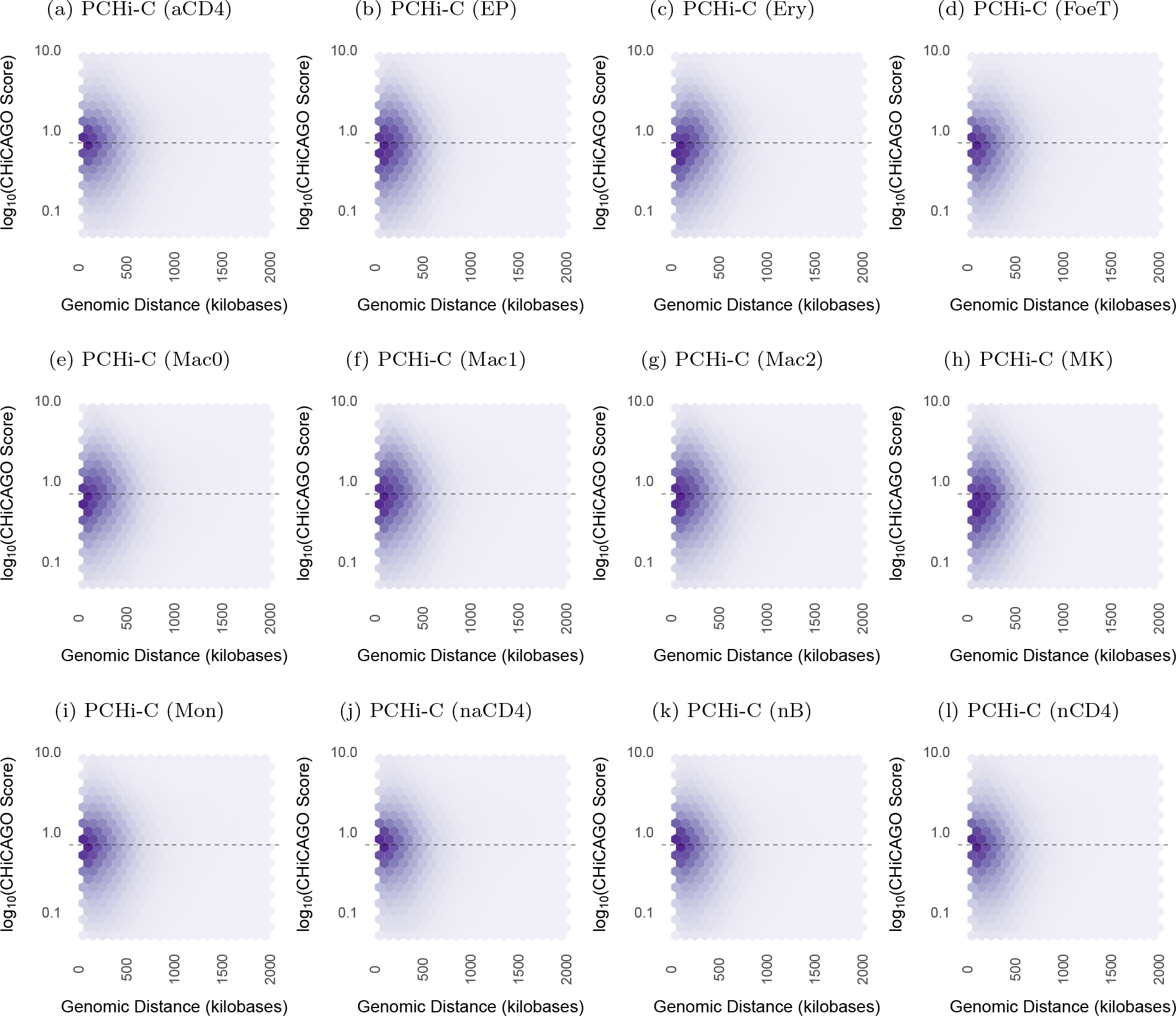
PCHi-C contact decay with genomic distance. Promoter Capture Hi-C contacts are anti-correlated with distance in 17 blood cell types. Decay is similar to Supplementary Figure 2, though here an interaction score (quantified by CHiCAGO [27]) is used rather than normalized contact frequency. A score above 5 denotes statistically significant interactions, and this threshold is indicated with a dashed line. Interaction distances are concentrated below 1mb, though a large number of significant interactions exist at 500kb, well beyond the 40kb dropoff in LD seen in Figure 3.

**Supplementary Figure 3:**
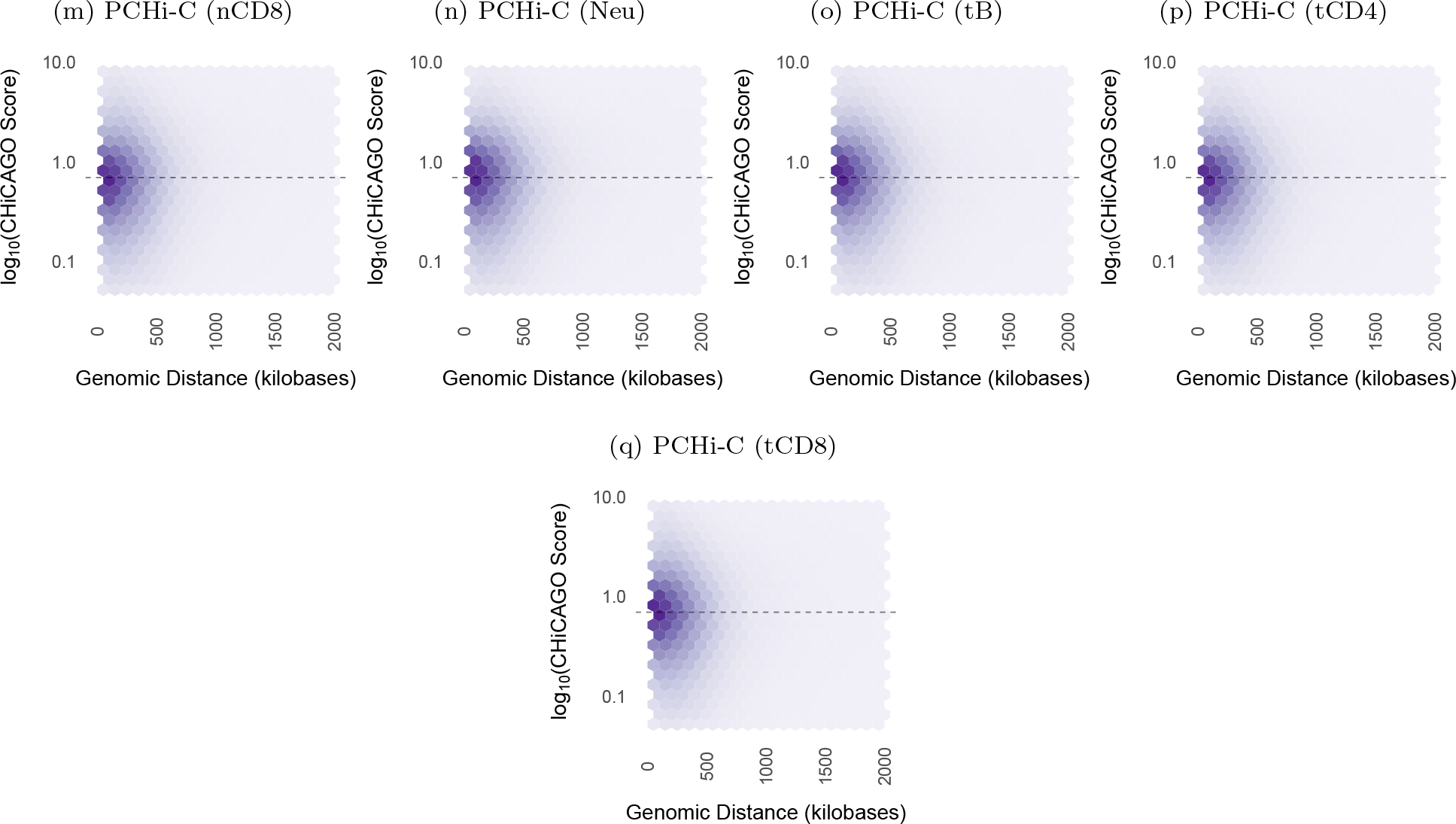
PCHi-C contact decay with genomic distance (cont.) Promoter Capture Hi-C contacts are anti-correlated with distance in 17 blood cell types. Decay is similar to Supplementary Figure 2, though here an interaction score (quantified by CHiCAGO [27]) is used rather than normalized contact frequency. A score above 5 denotes statistically signficant interactions, and this threshold is indicated with a dashed line. Interaction distances are concentrated below 1mb, though a large number of signficant interactions exist at 500kb, well beyond the 40kb dropoff in LD seen in Figure 3.

**Supplementary Figure 4:**
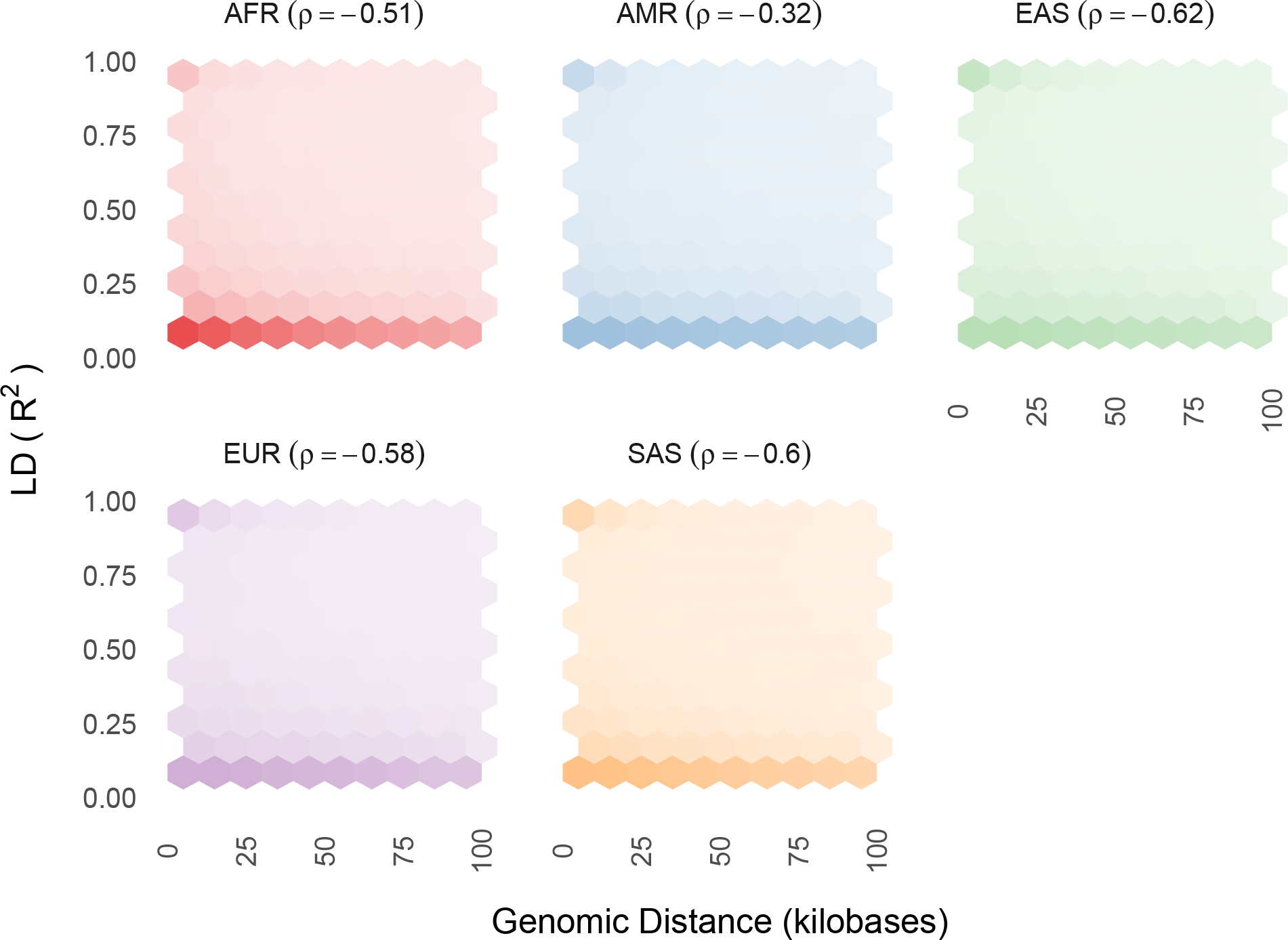
LD decay by superpopulation. Scaling of LD with genomic distance shows moderate anti-correlation for all superpopulations. Figure 2 shows combined LD scaling and observed Hi-C contact frequency scaling by cell line.

**Supplementary Figure 5:**
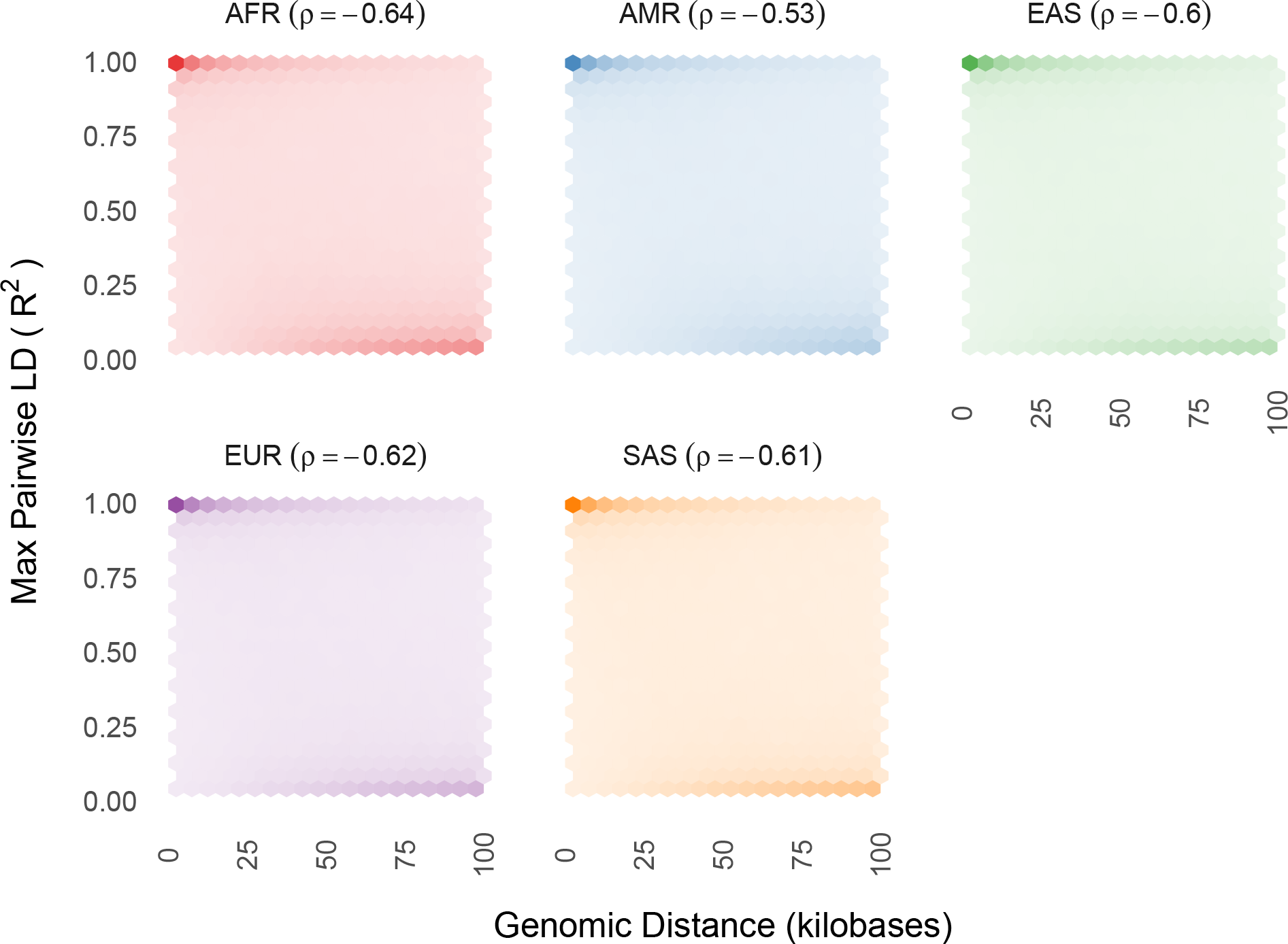
Interaction LD decay by superpopulation. Scaling of LD with genomic distance for SNPs located on statistically significant chromatin interactions shows moderate anti-correlation for all superpopulations. Figure 2 shows combined LD scaling (not restricted to interacting chromatin) and observed Hi-C contact frequency scaling by cell line.

**Supplementary Figure 6:**
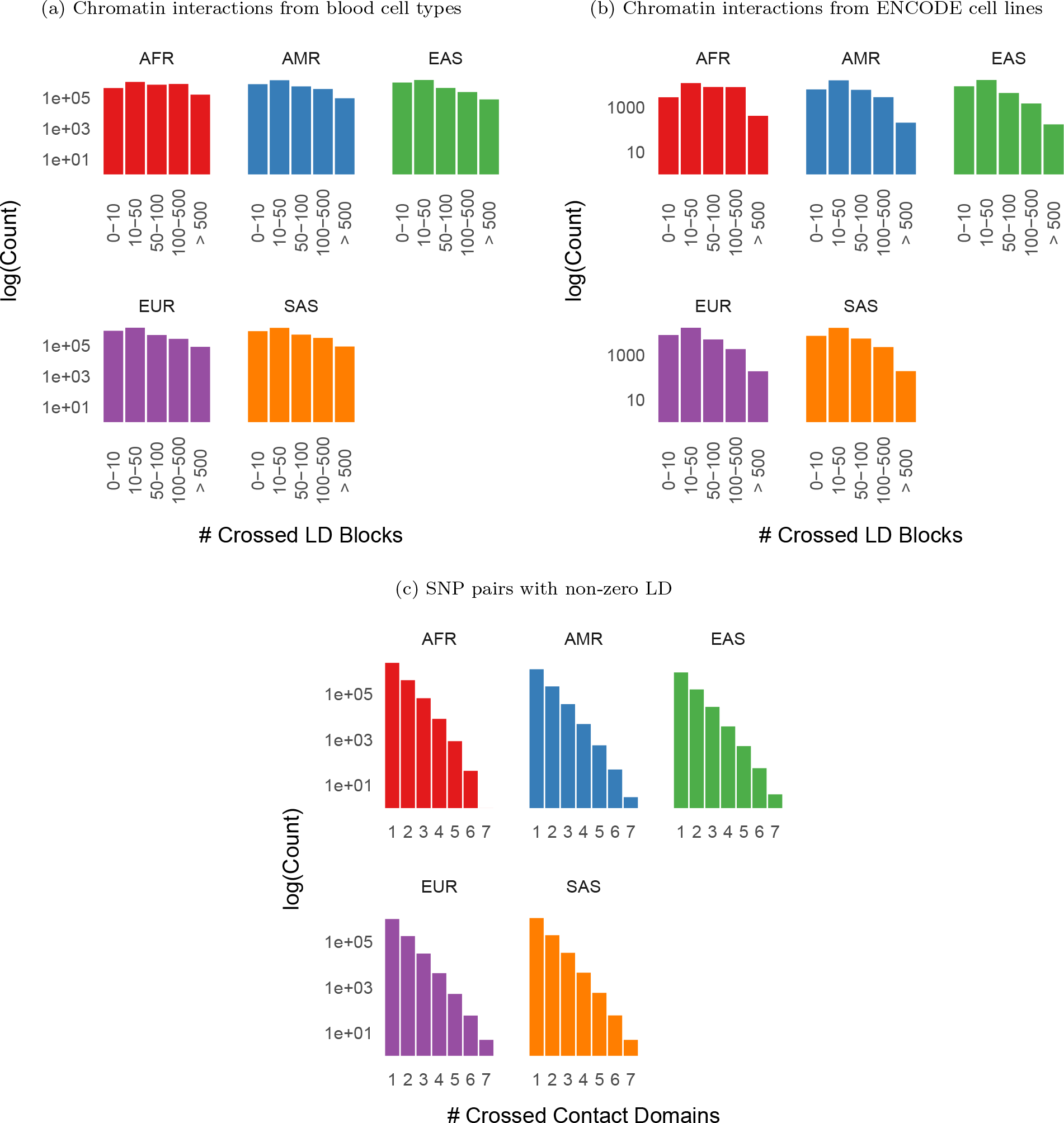
Interaction crossings. Number of LD blocks crossed by statistically significant chromatin interactions across all chromosomes in 17 primary blood cell types [12] (panel a) and 5 ENCODE cell lines [21] (panel b), grouped by superpopulation. Many statistically significant interactions cross hundreds or thousands of LD blocks. Panel c) shows the number of contact domains crossed by SNP pairs with non-zero LD, summed over 5 ENCODE cell lines, grouped by superpopulation.

